# BIOMAPP::CHIP: Large-Scale Motif Analysis

**DOI:** 10.1101/2023.11.06.565033

**Authors:** Jader M. Caldonazzo Garbelini, Danilo S. Sanches, Aurora T. Ramirez Pozo

## Abstract

**Background:** Discovery biological motifs plays a fundamental role in understanding regulatory mechanisms. Computationally, they can be efficiently represented as *kmers*, making the counting of these elEMents a critical aspect for ensuring not only the accuracy but also the efficiency of the analytical process. This is particularly useful in scenarios involving large data volumes, such as those generated by the *ChIP-seq* protocol. Against this backdrop, we introduce biomapp ::chip, a tool specifically designed to optimize the discovery of biological motifs in large data volumes.

**Results:** We conducted a comprehensive set of comparative tests with state-of-the-art algorithms. Our analyses revealed that biomapp ::chip outperforms existing approaches in various metrics, excelling both in terms of performance and accuracy. The tests demonstrated a higher detection rate of significant motifs and also greater agility in the execution of the algorithm. Furthermore, the smt component played a vital role in the system’s efficiency, proving to be both agile and accurate in *kmer* counting, which in turn improved the overall efficacy of our tool.

**Conclusion:** biomapp ::chip represent real advancements in the discovery of biological motifs, particularly in large data volume scenarios, offering a relevant alternative for the analysis of *ChIP-seq* data and have the potential to boost future research in the field. This software can be found at the following address: https://github.com/jadermcg/BIOMAPP-CHIP.

## 1 Background

*Motifs* are subsequences of length *k* that occur with high frequency in a set of sequences of dna, rna, or proteins. Formally, consider an alphabet Σ, which can be {*A, C, G, T }* for dna,{ *A, C, G, U}* for RNA, or a set of amino acids for proteins. A sequence *s* is an ordered list of symbols from Σand a *motif m* is a subsequence of *s* such that *m∈* Σ^*k*^, where *k* is the length of the *motif*. The discovery of motifs involves identifying these subsequences that frequently occur in a set of sequences, possibly with some variations [5] and [4].

It is important to note that *motifs* can be represented through *kmers* (contiguous subsequences of a given size k), making the efficient counting of these structures highly relevant. The representation of *motifs* as *kmers* allows for the simplification of complex computational problems, enabling efficient counting algorithms to be applied as a preliminary step in identifying biologically significant sequences [19].

In *motif* discovery, there are two main approaches: enumerative techniques and probabilistic techniques. Both have their own advantages and disadvantages, and the choice between them depends on the specific needs of the analysis at hand. Enumerative techniques, as the name suggests, aim to identify *motifs* by exhaustively enumerating all possible subsequences within a set of sequences. These methods are generally more accurate as they consider all possibilities. However, they are computationally intensive and may not be feasible for large volumes of data or for *motifs* of significant length [10].

On the other hand, probabilistic techniques such as *Gibbs Sampling* (gs) and *Expectation Maximization* (em) employ statistical models to estimate the presence of *motifs*. These methods are generally faster in handling large volumes of data; however, they are sensitive to initial conditions and model parameters, which may compromise accuracy. Both approaches have their merits: while enumerative techniques are typically more accurate, probabilistic techniques are more time-efficient. Nevertheless, it is possible to integrate these two approaches to leverage the advantages of both. For example, one could use an enumerative approach to filter candidates and a probabilistic approach for final optimization [16].

This interconnection between *kmer* counting and *motif* discovery highlights the importance of efficient algorithms in both domains for the acceleration and enhancement of bioinformatics analyses. In this context, algorithms for *kmer* counting serve as initial tools in the processing chain for discovering biological *motifs*, especially in analyses that involve large volumes of data, such as *ChIP-seq* (Chromatin Immunoprecipitation followed by Sequencing) data.

The *ChIP-seq* (Chromatin Immunoprecipitation followed by Sequencing) protocol is a modern technology that allows the identification of dna-protein interactions on a genomic scale. This method has become an indispensable tool in the discovery of biological *motifs*, as it provides a comprehensive mapping of protein-binding regions throughout the genome. One of the major challenges of *ChIP-seq* is the substantial volume of data generated. The handling, storage, and analysis of such data require robust computational infrastructure and efficient algorithms, as traditional *motif* discovery techniques may not be suitable for such a magnitude of data [9].

In addition to the data volume challenge, complexity and the presence of noise are also significant factors. The *ChIP-seq* protocol is susceptible to various types of artifacts and noise that can introduce errors in *motif* identification. Therefore, robust preprocessing and filtering methods are required before the *motif* discovery stage itself. Many current *motif* discovery algorithms were not designed to handle the challenges imposed by *ChIP-seq*, such as large data volumes and complexity in sequence structure. This often results in a trade-off between accuracy and computational efficiency [12].

In this context, biomapp ::chip (*Biological Application for ChIP-seq data*) is designed to address these challenges. Our algorithm adopts a two-step approach for *motif* discovery: counting and optimization. In the counting phase, the smt (*Sparse Motif Tree*) is employed for efficient *kmer* counting, enabling rapid and precise analysis. For the optimization stage, biomapp ::chip employs an enhanced version of the EM algorithm, aimed at improving accuracy in *motif* identification.

## 2 ImplEMentation

Implemented in C++ and R, biomapp ::chipX) combines analytical and numerical methods optimized for in-depth data treatment, particularly data derived from *ChIPseq* experiments. The objective of this approach is to build effective solutions that assist researchers and experts in the field of molecular biology, more specifically in the study of conserved sequences, thereby facilitating complex investigations that involve sequence motif analysis. Through the appropriate combination of methods, some already established and others specifically created for our framework, biomapp ::chip offers a robust and adaptable tool for the scenario of functional genomics. Figure 1 illustrates the general pipeline, all the framework modules, and their respective interactions.

**Fig. 1.**
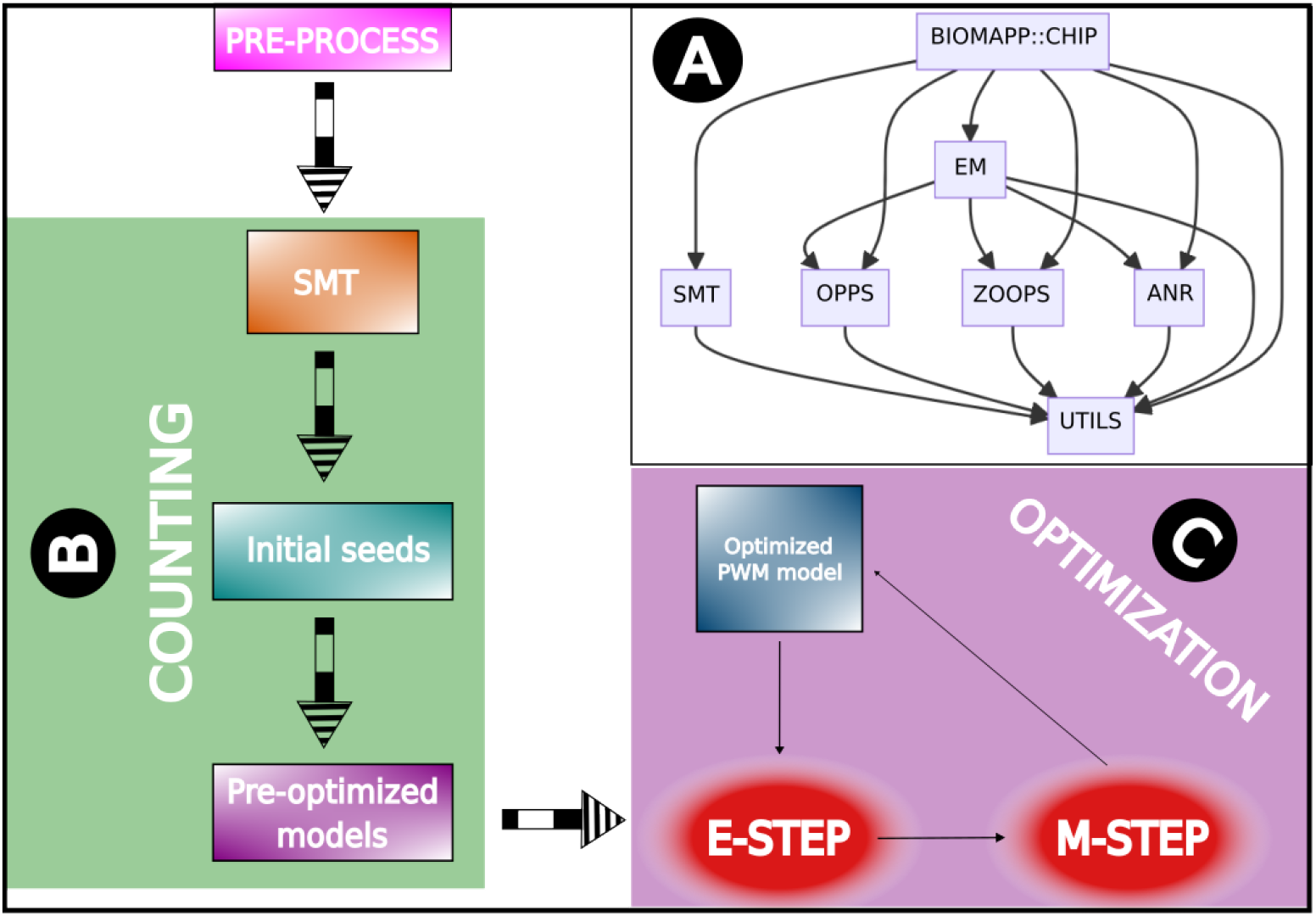
BIOMAPP::CHIP framework pipeline. (A) The main workflow is structured in various interconnected modules. Each module is responsible for a specific part of the overall functionality of the package. Preprocessing involves identifying and normalizing peaks in the data. (B) In the counting step, the dataset is loaded, a background model is generated, followed by the construction of the SMT. Enriched *kmers* are extracted and initial models are created. (C) Optimization occurs by refining the seeds using the FAST-EM algorithm.

As illustrated in Figure 1, the framework initiates its process with the preprocessing of the input sequences. These are then forwarded to the SMT, responsible for the efficient counting of *kmers*, generating seeds that have already gone through an initial stage of optimization as a result. These seeds are subsequently refined by the fast-em algorithm. The architecture is modular and composed of seven main modules: utils, smt, oops, zoops, anr, em, and biomapp ::chip. The UTILS module acts as the backbone of the system, providing essential functionalities used by all the other modules.

The smt module is developed on the foundation provided by the utils, and the oops, zoops, anr, and em modules are likewise constructed on this foundation. In addition to being supported by the utils, the EM module has interdependencies with the oops, zoops, and anr modules. The biomapp ::chip module, in turn, represents the highest level of complexity as it integrates the functionalities of all the previous modules to execute its operations. Parallelization is supported in all modules through the openmp (https://www.openmp.org/) and thread building blocks (https://www.intel.com/content/www/us/en/developer/tools/oneapi/onetbb.html) libraries. The source code of biomapp ::chip is available at: https://github.com/jadermcg/biomapp.

### 2.1 Preprocessing

Before delving into the method description, it is important to understand how the data are acquired. They can be obtained from various public and private sources. For more information on the *ChIP-seq* databases used in this work, please refer to Section 3.The data acquisition process can be summarized in the following steps:

1. Experiment execution: the proteins of interest are immunoprecipitated along with the associated dna fragments.
2. Raw data processing: the raw sequencer data are processed to obtain short dna sequences, called reads.
3. Peak identification: specific programs, such as macs [22] or sicer [21], are used to identify regions of reads enrichment, called peaks.
4. Pre-processing: removal of peaks in non-specific regions, peak filtering based on quality criteria, data normalization, and removal of spurious peaks.
5. Analysis: performing the analysis of interest, which may include: motif extraction, functional annotation, regulatory network analysis, conservation analysis, among others.

The steps 1 and 2 are part of the biological experiment and are generally conducted by a biologist. Step 3, called enrichment analysis, aims to identify the peak regions, ranging from 100 to 300 bp^1^, where there is a high probability that the fragments of interest are present. Step 4 is critical as it involves the removal of repetitive and low-complexity sequences. The final step consists of the analysis of the pre-processed data.

The biomapp ::chipX) starts operating at the end of step 3, where it receives as input the peak regions properly extracted from the genome. In step 4, specialized algorithms are used for pre-processing, and step 5 is fully implemented. The biomapp ::chip consists of three main phases: pre-processing, initialization, and optimization.

The pre-processing phase involves adapting the dataset for the execution of experiments, such as standardizing the size of the sequences and converting all characters to uppercase. However, the main goal of this stage is to identify low-complexity sequences, transposable elements such as alus (*Arthrobacter luteus*) and sines (*Short Interspersed Nuclear Elements*), *E. coli* insertions, or any other components that could lead to inaccurate conclusions in subsequent steps. This is carried out with the help of specialized algorithms like dust [20] and repeat masker [18]. This phase involves the elimination of redundant sequences and error correction, ensuring the quality and integrity of the data.

### 2.2 Counting

The counting phase takes as input pre-processed peak regions, where it is presumed that there are fragments from the distribution of interest surrounded by background distribution elements. The purpose of the initialization algorithm is to efficiently identify, count, and group enriched *kmers*, as well as to perform hypothesis tests to determine their statistical significance. The core of this stage consists of two components specifically developed for this purpose: i) smt: a scalable data structure based on suffix trees responsible for storing all *kmers* from the main dataset; ii) kdive: an efficient search algorithm inspired by depth-limited search and the branch and bound algorithm.

The counting phase follows a set of steps to identify enriched *kmers* in the main dataset and generate initial models based on this information. First, the algorithm loads the pre-processed data, which contains the nucleotide sequences of interest. Next, if available, control sequences are loaded. Otherwise, they are generated by shuffling the main dataset using the Markov method [6] or the Euler method [1]. A background probabilistic model is then generated from the control data, which will serve as a reference for identifying enriched *kmers*. The value of k is then estimated, or a predefined value is used, depending on the approach adopted. Based on the peak regions, the smt data structure is built, allowing the efficient extraction of enriched *kmers*.

### 2.2.1 SMT

The smt is represented by a two-dimensional data structure *M*^*ν×*6^, where *ν* is the number of nodes, implemented from fixed-width text fragments. Unlike ktrees or strees, the smt is specifically designed to deal with *kmers*. It is lightly inspired by room squares theory [2], in which each element *M*_*i,j*_ is either empty or has a value set between 1 and *ν*. Its main goal is to efficiently store all *kmers* belonging to the main dataset, along with their representation and count, allowing for rapid searches. Thus, each row of *M* represents a node {1, 2, 3, …, *ν}*, columns 1 to 4 represent the nucleotides, and the last two columns, 5 and 6, represent the number of times a fragment appeared in the dataset and its respective address or numeric representation.

For example, if *M*_1,*T*_ = 10, then the symbol T is present between nodes 1 and 10. In comparison to similar structures, smt is more efficient for storing and searching fixed-size fragments, and its construction has linear time complexity *O*(*k*), where *k* is the size of the sequence of interest. Furthermore, it is possible to execute various algorithms on the smt in linear time, such as exact search and approximate search, which, for example, allow for the rapid retrieval of the most abundant *kmers* in the dataset. Algorithm 1 shows how the smt is constructed.

#### Algorithm 1

create-SMT

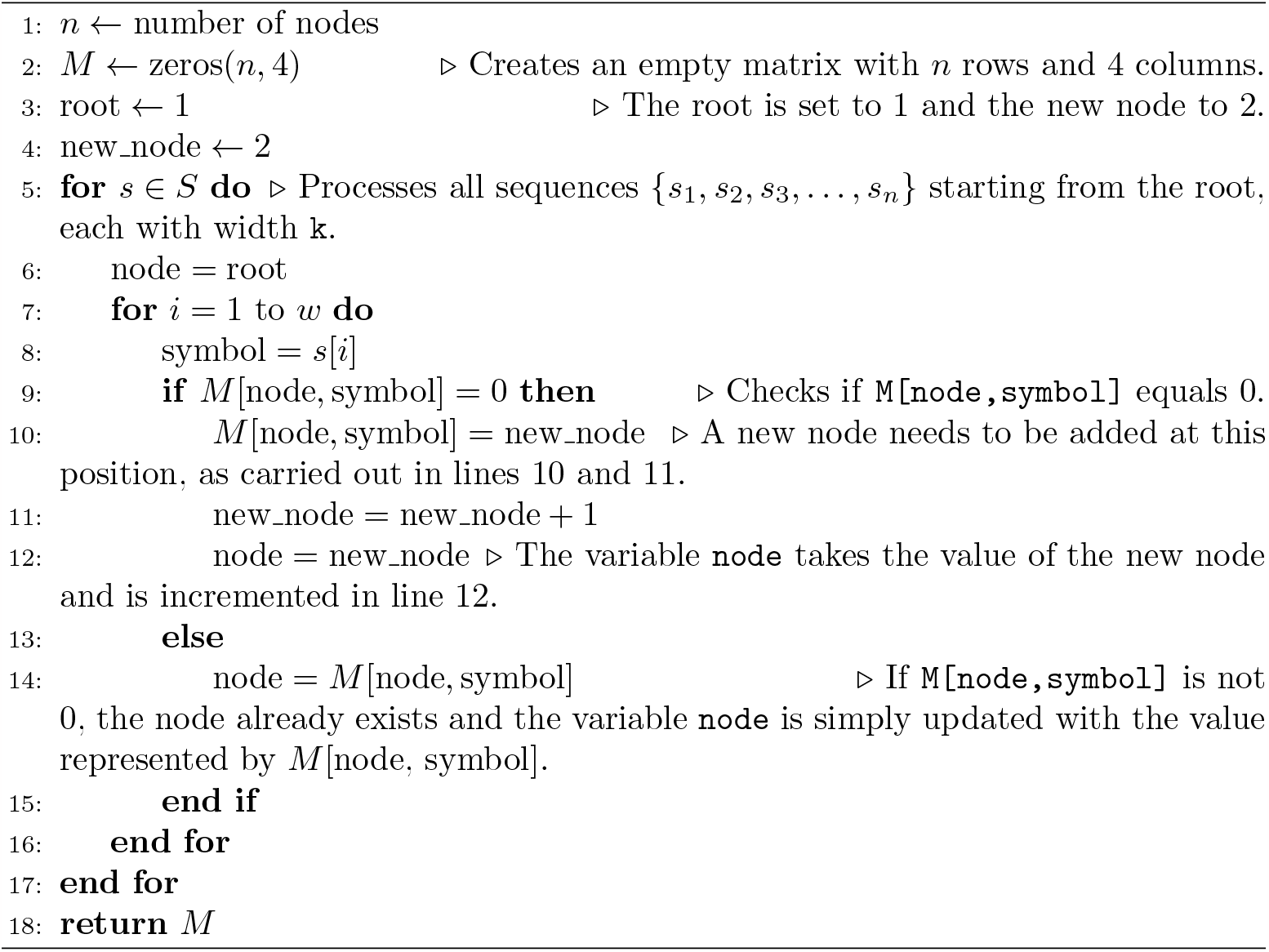

In line 3, the root (root) is set to the value 1 and the new node (new node) as 2. In line 5, all the sequences {*s*_1_, *s*_2_, *s*_3_, …, *s*_*n }*_ are processed starting from the root, each of which has a width of k. In line 9, the algorithm checks if M[node, symbol] is equal to 0. If this occurs, it means that there is no edge with the symbol defined by the variable `symbol` leaving from the variable `node` and going towards another node. In other words, it is necessary to add a new node at this position, which is done in lines 10 and 11. Finally, the variable node takes the value of the new node `(new node)` and in line 12, the new node is incremented. In line 13, if `M[node, symbol]` is different from 0, then the node already exists, and in line 14, the variable node is simply updated with the value represented by *M* [node, symbol].

The space complexity of smt in the worst case is *O*(*σν*), where *σ* is the number of symbols in the alphabet, in this case 4, and *ν* is the number of nodes in the tree. The value of *ν* directly depends on the size and variance of the fragments. Thus, the higher the variance, the greater the number of nodes. An inherent characteristic of smt is that most elements in `M` remain empty. This behavior is recurrent and can be calculated. In particular, we can compute the expected occupancy level of M. To do so, consider that the maximum and minimum number of children a node can have is 4 and 0, respectively. The average would then be 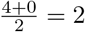children per node.

Your inclusion of the *ω* parameter to represent the occupancy level of the matrix *M* provides a nuanced understanding of real-world performance. This is particularly relevant because computational structures rarely operate at their theoretical worst-case limits in practical applications. By calculating the average-case scenario using *ω*, you not only present a more realistic model but also offer an insightful optimization strategy. Specifically, the fact that *O*(*ων*) *< O*(*σν*) implies that under average conditions, the data structure is more space-efficient than one might initially surmise from the worst-case scenario alone.

Thus, we can expect that only half of the capacity of `M` will be utilized, which characterizes a significant waste of space. For this reason, the create-SMT algorithm was implemented using a high-performance sparse matrix through the armadillo linear algebra library [17], employing the C++ programming language. The time complexity of smt is simpler to calculate and in all cases is *O*(*mk*), where m represents the quantity and k the width of the fragments. To see this, we only need to consider that each fragment is composed of k symbols. Since there are m fragments, mk nucleotides will need to be processed in total. However, if we consider that *k*≤20 and that m can grow indefinitely, then we have *k*≪*m*. As k has a constant upper limit, we can approximate the complexity to *O*(*mc*) = *O*(*m*).

#### 2.2.2 KDIVE

While kdive stands as the most important algorithm for analyzing conserved sequences in the context of SMT, it is not the only option available. Other algorithms that operate on the smt include KSEARCH, IUPACSEARCH, and KHMAP. Each of these methods offers distinct advantages and functionalities, and they are discussed in detail in the *Supplementary data*.

KDIVE was created with the objective of performing efficient text fragment searches in the SMT, even if these present up to d degenerations. In other words, kdive should return true for the search even if the query string contains up to d mismatches. It is important to highlight that the algorithm assumes as a premise that the probability of a MISMATCH occurring is the same for any position in the sequence.

For example, if the probability of a match is equal to 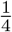, then that of a MISMATCH is 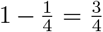. Therefore, the occurrence of a MISMATCH is three times more likely than that of a match. This characteristic is important because it alters the expected number of computations carried out and consequently modifies the algorithm’s complexity. The construction of kdive was based on limited depth search [15] and the Algorithm (branch and bound). Algorithm 2 shows the basic functioning of KDIVE.

##### Algorithm 2

KDIVE

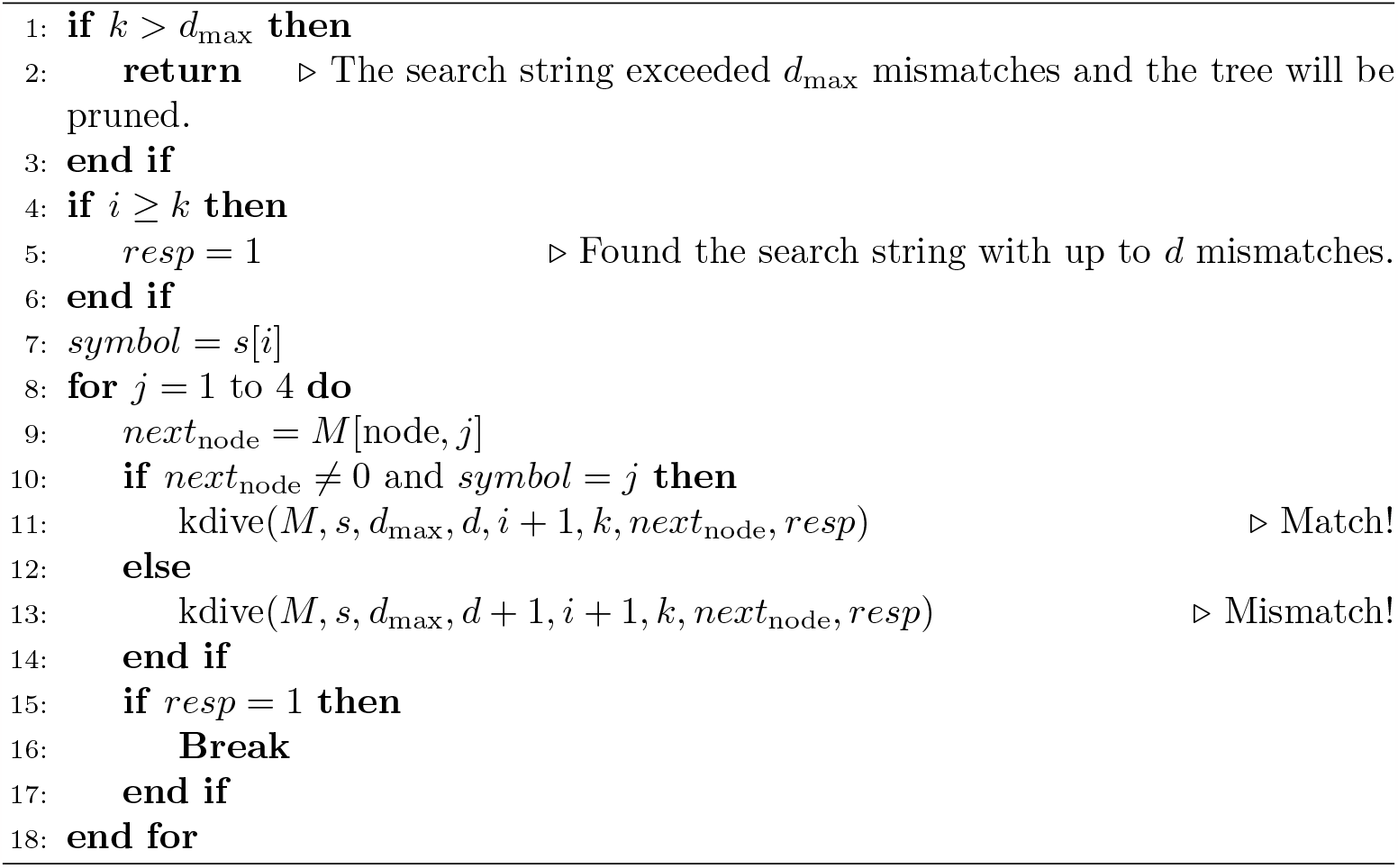

Algorithm 2 was implemented recursively and takes as its main parameters the smt matrix, the fragment `s`, and the total number of allowed degenerations d. If *s* ∈ *M* with at most `d` mismatches, the algorithm returns the boolean value true, indicating the presence of the fragment with up to d mutations.

The algorithm performs its task by traversing the smt to identify matches between the search string and the stored *kmers*, taking into account the number of allowed mismatches. It is worth noting that this algorithm can be easily adapted to return the complete set of degenerate elements, instead of just a boolean value (true or false). This modification can be useful in certain applications, as it allows for more detailed information about the matches found. For example, it is possible to identify which *kmers* in the smt correspond to the search fragment, considering the allowed mutations.

The parameters *d*_max_, d, i, and k allow for the control of searching for exact or approximate matches. The recursion enables the algorithm to traverse the tree in depth and check all possible paths until it finds a match or reaches the maximum depth defined by the parameter *d*_max_. A possible call to Algorithm 2 could be kdive(M, s, *d*_max_ = 2, d = 0, i = 1, k = 10, node = root, resp = false). In this case, the algorithm expects to find a match even if up to *d* = 2 mutations are detected in sequences of size *k* = 10.

Algorithm 2 has two base cases, shown on lines 1 and 4. The first checks if the number of mutations has exceeded the limit *d*_max_, that is, if *d≥d*_max_. If this condition is met, a pruning will occur in the search. The second base case checks if the algorithm has completely analyzed the query string, which will happen if *i≥k*. If this condition is true, then the pattern has been found and the *resp* variable is updated to the value TRUE.

The time complexity of kdive can be measured through the Negative Binomial distribution. Consider a smt with *ν* nodes, *d* mismatches, and fragments of width *k*. Also consider the random variable *X∼NB*(*d, p*), which counts the number of comparisons that the kdive algorithm performs. It’s easy to verify that 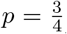, since if the probability of a match is 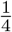, the probability of a MISMATCH will be 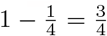 therefore 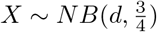. The PMF *P*_*x*_(*d, p*) is given by Equation 1 and the expectation by Equation 2.

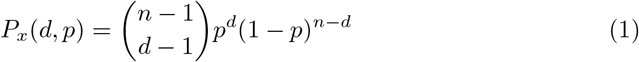

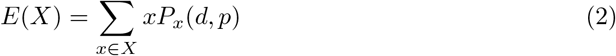

The Negative Binomial distribution is a generalization of the geometric distribution, which is defined as *X ~ GEOM* (*p*) = *NB*(1, *p*). Therefore, the geometric distribution is the Negative Binomial distribution with *d* = 1, making it possible to use it for calculating the expected value. For example, consider *X∼NB*(1, *p*), *Y ~ NB*(1, *p*), and *Z* = *X* + *Y*. If *X* and *Y* are independent, then *Z* = *X* + *Y∼NB*(2, *p*). Therefore, we can calculate 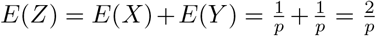. In this way, *X ∼ NB*(*d, p*) can be written as *Z* = *X*_1_ + *X*_2_ + *X*_3_ + *…* + *X*_*d*_, where *X*_*i*_ *∼ NB*(1, *p*). Calculating 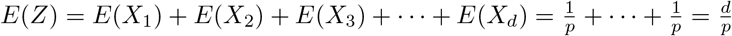.

With this result, we can write that the time complexity of kdive is 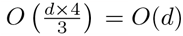. It is easy to verify that the larger the size of *d*, the more operations will be necessary. In particular, the complexity grows linearly with *d*. It is important to highlight that MOTIFS rarely exceed a width of 20 nucleotides, and the number of mutations is generally no more than 20% of their size. Considering this, we can expect to find an average of *d* = 5 mismatches, which results in 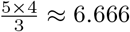 comparisons per fragment.

We can calculate the complexity of the kdive algorithm in relation to the size of the input sequences. Consider a dataset with *n* sequences of width *t*. We have a total of *m* = *t−k* + 1 fragments of size *k* in each sequence of size *t*. Let *X* be the number of comparisons made for each fragment. We know that *X* follows a negative binomial distribution with parameters 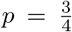 and *d*, and that 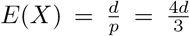. Therefore, the number of comparisons per sequence will be 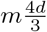. If we consider, for example, a fragment of size *k* = 20 with *d* = 5 mutations, then 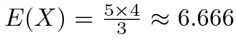, which will result, on average, in 6.666 … *×m* = *O*(*m*). It is interesting to highlight that this value will be even lower in practice because kdive uses the branch and bound algorithm to prune the search tree.

Even in the worst-case scenario, the algorithm will still exhibit linear asymptotic behavior. In a less detailed analysis, we arrive at the following complexity: *k*(*m−k* + 1) = *km−k*^2^ + *k*, in which the dominant term is *k*2, and therefore the complexity of this algorithm will be of the order of *O*(*k*^2^). However, *k* is often much smaller than *m*, and in this way we can make a more refined analysis. Given that *k*≪*m*, the expression *k*(*m−k* + 1) can be approximated as: *k*(*m − k* + 1)*≈km*.

The complexity of the algorithm is generally dominated by the term *km* and can be expressed as *O*(*km*). However, considering that *k* is not greater than 20 and *m* can grow indefinitely, we can adjust the complexity analysis. As *k* has a constant upper limit (*k*≤20), the complexity of the algorithm is mainly influenced by the term *m*. Therefore, we can express the complexity of the algorithm as *O*(*m*).

#### 2.2.3 Optimization

The aim of this stage is to optimize the initial seeds obtained in the previous phase through the em algorithm. The use of this algorithm allowed refining the parameters associated with the seeds, making thEM more accurate representations. This optimization process is important for the quality of the resulting models and for the success of subsequent analyses.

In this phase, pre-optimized seeds obtained from the smt are used as a starting point for running the em algorithm. The em algorithm is responsible for refining these seeds, adjusting their parameters in order to maximize the likelihood of the given observations. As a result of this phase, each seed is transformed into a pwm model, which represents a more accurate and adjusted description of the motif in question. This pwm model can then be employed on new data to find patterns that have not yet been labeled.

To select the appropriate variant of the em algorithm for *ChIP-seq* data, it’s important to consider the protocol’s nuances, which include the presence of noise and non-specific signals. While *ChIP-seq* aims to capture sequences with the motif of interest, not all peaks may contain it due to various factors. Therefore, the zoops model is generally the most fitting for motif analysis in *ChIP-seq* data, allowing for the motif’s occasional absence. Depending on the biological context, anr or oops models may also be relevant.

#### 2.2.4 FAST-EM

The fast-EM represents a significant optimization over traditional oops and zoops models. This algorithm was inspired by the implementation provided by [7] [8], whose modification not only speeds up the execution time but also enhances efficiency in handling larger data sets. In other words, fast-EM enhances the capabilities of the original models, making them more adaptable and robust when faced with large volumes of information.

The calculation of marginal probabilities represents one of the most computationally demanding phases in the em algorithm. This stage requires computing the probability of a *kmer* being found at each of the *n×*(*t−k* + 1) valid positions, given the *n* input sequences. This process can become a significant bottleneck, especially in *ChIP-seq* experiments where a large volume of sequences is often dealt with, in turn making the value of *n* considerably high. Although it is possible to mitigate this computational challenge by using only a subset of the sequences as a sample, this approach compromises the predictive power of the model.

To understand its workings, we need to recap some concepts. The bayes’ rule for the em algorithm states: 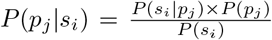, which can be interpreted as: “the probability of a motif existing at position *p*_*j*_ given we are observing sequence *i*, divided by the marginal probability of *s*_*i*_.” The marginal probability can be written as the weighted sum: *p*(*s*_*i*_) = *P* (*s*_*i*|_*p*_1_)*P* (*p*_1_) + …+ *P* (*s*_*i* |_*p*_*m*_)*P* (*p*_*m*_). If we consider that a MOTIF can be in any position with equal likelihood, then this sum simplifies to: *p*(*s*_*i*_) = *P* (*s*_*i*_ *p*_1_) +… + *P* (*s*_*i*_ *p*_*m*_).

To compute each term of the marginal probability, it is necessary to employ both the positive model (*α*) and the negative model (*β*) through the following equation: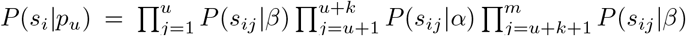, for 1 *≤ u ≤ m*. In other words, this equation must be run for all *m* valid positions in each sequence. The FAST-EM algorithm does this by computing 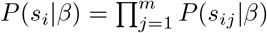 just once. Then, for each 1 *≤ u ≤ m*, this value is divided by 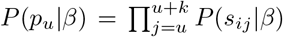and multi-plied by 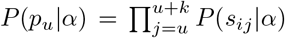. To work on a logarithmic scale, simply replace multiplications with additions and divisions with subtractions. This simple optimization, when combined with multithreading techniques, makes the fast-EM algorithm significantly faster and, therefore, better suited for handling large volumes of data.

## 3 Results and discussion

In this section, we will show and analyze the results obtained by the biomapp-chip framework in comparison with state-of-the-art algorithms. The biomapp-chip was rigorously compared with state-of-the-art algorithms, including meme [3], PROSAMPLER [13], and homer [11]. In the experiments conducted, two distinct types of data were employed to evaluate the effectiveness of our method. Synthetic data were used for load tests, allowing a comparative analysis of time consumption and RAM memory usage between the different approaches. On the other hand, real data were employed to assess accuracy, thus ensuring a more realistic evaluation of its applicability in practical scenarios.

In addition to comparisons with state-of-the-art methods, we also conducted a direct evaluation between the smt and jellyfish [14], a widely-used tool for *kmer* counting. The aim was to understand the efficacy of the smt algorithm in relation to established methods in the literature. Details on this comparison, including performance metrics, are provided in *SupplEMentary data*. This additional analysis reinforces the robustness of our approach and offers further insights into the performance of the smt algorithm.

It is important to emphasize that all performance metrics presented in this study have been rigorously evaluated. Load tests, which include measurements of both execution time and memory consumption, were standardized and quantified with precision. Execution time was measured in seconds, while memory consumption was recorded in megabytes, thereby offering a transparent and comparable evaluation of the algorithm’s performance. In addition to performance metrics, the accuracy was assessed using various distance and correlation measures as benchmarks. These included euclidean distance, manhattan distance, hellinger distance, pearson correlation, bhattacharyya coefficient, and sandelin-wasserman similarity. These measures were employed to rigorously evaluate the model against reference standards.

## 3.1 Synthetic data

The generation of synthetic data was carefully planned to allow a comprehensive comparison between biomapp-chip framework in relation to other motif discovery algorithms. For the comparison, 50 datasets were generated, with sizes ranging from 1*×* 10^5^ to 2*×* 10^7^ bases. Each of these datasets was submitted to all algorithms with k-mer sizes (k) ranging from 5 to 30 bases. This experimental design allowed for a rigorous evaluation of performance and efficiency in k-mer counting under different load and complexity conditions.

### 3.2 Real data

Real data were extracted from the jaspar database version 2022 along with the genomic position files (.BED). These experiments form a fundamental step in validating our approach. By using real data, we can evaluate the performance of the algorithms in practical applications, increasing the reliability of the results and the robustness of our conclusions. Table 1 presents a detailed analysis of the data volume (measured by the number of peak regions) generated by different types of experiments available in the jaspar 2022 database.

**Table 1.** Number of peak regions extracted from various experiments available in the jaspar 2022 database. It is clearly noted that the *ChIP-seq* protocol is responsible for generating the most significant volume of data.

|  | TYPE | MIN | MAX | AVG |
| --- | --- | --- | --- | --- |
| 1 | ChIP-seq | 22 | 322803 | 17538.78 |
| 2 | CAP-SELEX | 206 | 22850 | 9878.26 |
| 3 | HT-SELEX | 55 | 91919 | 9856.47 |
| 4 | NA | 84 | 105198 | 8801.62 |
| 5 | NCAP-SELEX | 2249 | 13498 | 6792.00 |
| 6 | PBM | 98 | 100001 | 4424.22 |
| 7 | COMPILED | 6 | 28379 | 3661.81 |
| 8 | DAP-seq | 22 | 19758 | 1222.74 |
| 9 | SMiLE-seq | 998 | 1001 | 999.97 |
| 10 | ChIP-exo | 48 | 1000 | 568.80 |
| 11 | SELEX | 9 | 1001 | 561.11 |
| 12 | ChIP-chip | 71 | 4737 | 475.80 |
| 13 | SELEX-seq | 384 | 384 | 384.00 |
| 14 | Universal protein binding microarray (PBM) | 99 | 101 | 100.00 |
| 15 | PBM, CSA and/or DIP-chip | 8 | 102 | 99.12 |
| 16 | EMSA | 29 | 100 | 64.50 |
| 17 | Bacterial 1-hybrid | 13 | 60 | 23.13 |
| 18 | DNaseI footprinting | 10 | 41 | 16.33 |

The *ChIP-seq* protocol stands out as the main source of data, generating the largest volume of information, with an average number of peak regions of 17,538.78. The data volume generated by this protocol varies considerably, ranging from a minimum of 22 to a maximum of 322,803 peak regions. Other significant protocols in terms of data volume include cap-selex and ht-selex, which generated an average of 9,878.26 and 9,856.47 peak regions, respectively. Although these protocols generate considerable data volumes, they are still significantly below *ChIP-seq*.

In the experiments involving real data, datasets containing more than 10,000 sequences were selected. This criterion resulted in an initial set of 148 distinct datasets. However, 17 of these datasets were subsequently excluded from the analysis, as they were still in the process of validation. Therefore, the final set for evaluation consisted of 131 validated datasets. The choice of this selection criterion aims to ensure a sufficiently large and varied sample to rigorously and comprehensively evaluate the performance of the algorithms. This approach allows not only to test the efficacy of the algorithms on real data, but also provides an in-depth analysis of their applicability in scenarios closer to the experimental conditions frequently encountered in research in the area of biological motif discovery.

### 3.3 Parameters

In order to ensure the reproducibility of experiments, we established the parameters for each algorithm according to specific guidelines. The standardization of these settings is important to guarantee the integrity and comparability of the results obtained. Furthermore, all experiments were run on Intel Xeon CPU E5-2673 v4 2.30GHz servers with 8Gb of ram memory with linux/ubuntu 22.04 operating system. The parameters used in the experiments followed those displayed in Table 2.

**Table 2.**
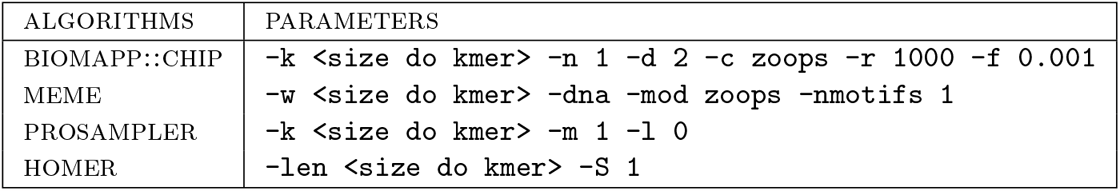
Parameters configured for the algorithms used in the research. Each algorithm is listed alongside its respective parameters to ensure the reproducibility of the experiments.

It is essential to clarify some aspects regarding the choice of parameters. All methods were adjusted to identify only the most effective model, as evidenced by the options `-n 1, -nmotifs 1, -m 1`, and `-S 1` for the smt, meme, prosampler, and HOMER algorithms, respectively. The ZOOPS model was employed in the biomapp ::chip and meme algorithms, while homer uses the zoops model intrinsically. The prosampler algorithm, on the other hand, does not offer the option of selecting this model. Additionally, the parameters `-r 1000` and `-f 0.001` are used to govern the convergence of biomapp ::chip. The iterative process will be halted if the increment in the score difference between two successive iterations is below -f 0.001. Otherwise, the algorithm will continue until it reaches a limit of -r 1000 iterations.

#### 3.3.1 BIOMAPP-CHIP on synthetic data

The aim of these experiments was to measure the performance of the biomapp ::chip framework on a diverse set of synthetic datasets. These data were generated with variable Parameters to simulate different scenarios and complexities associated with the real world. The use of synthetic data offers a controlled platform that allows isolating and evaluating the performance and efficacy of algorithms under well-defined conditions. This analysis procedure is essential for the initial validation of the adopted methodological approach, serving as a preliminary step before handling real data.

These trials were designed to collect basic data on the time consumption and RAM memory usage by the biomapp ::chip algorithm. This information was compared with time and space metrics obtained from other established algorithms in the field, such as meme, prosampler, and homer. The purpose of this comparison was to understand how biomapp ::chip stands in relation to other solutions and identify the strengths and weaknesses of each approach.

To carry out the experiments, the *k* Parameter, representing the size of the *kmers*, was systematically varied within a range that covers values from 5 to 30, thus keeping\ in line with the criteria established in the smt load tests. In this way, 5200 experiments were performed, 1300 for each algorithm. The capture of relevant metrics, which include both time consumption and RAM memory allocation, was conducted using the unix/linux **/usr/bin/time -v** command, ensuring consistent and comparable data collection across all algorithms under study.

Figure 2 illustrates the global averages of the tests for each algorithm, considering both time and space consumption. biomapp ::chip proved superior in terms of time efficiency, closely followed by meme. On the other hand, PROSAMPLER and HOMER showed less efficacy in this aspect, requiring longer periods for execution. Regarding memory usage, meme led in performance but was closely followed by biomapp ::chip. The latter employs the smt data structure to initialize its models, which demonstrates the efficiency of this structure in terms of memory usage. As observed in the time dimension, HOMER and PROSAMPLER occupied lower positions in memory consumption.

**Fig. 2.**
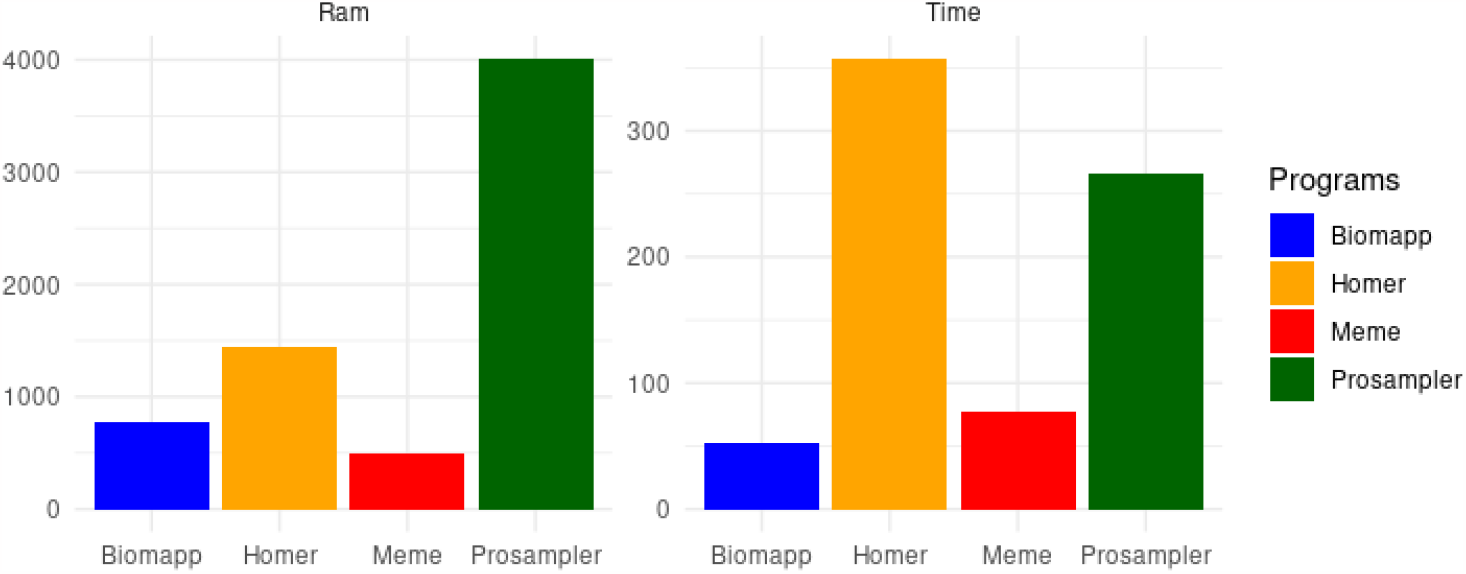
Performance comparison between the algorithms biomapp-chip, meme, homer, and prosampler, focusing on time (measured in seconds) and memory (measured in Mbytes) consumption. meme stood out for efficient memory usage, while BIOMAPP-CHIP was the fastest in terms of execution time.

The Figure 3 analyzes the execution time of the four algorithms – biomapp ::chip, meme, homer, and prosampler – as a function of the k size. It is noticeable that biomapp ::chip is the most time-efficient algorithm, closely followed by meme, both maintaining an average execution time below 100 seconds for all evaluated cases. In contrast, the average execution times for prosampler and HOMER were substantially higher, with prosampler exceeding 200 seconds and homer surpassing 400 seconds. Additionally, the analysis also reveals that the execution time for all evaluated algorithms increases as the size of k grows.

**Fig. 3.**
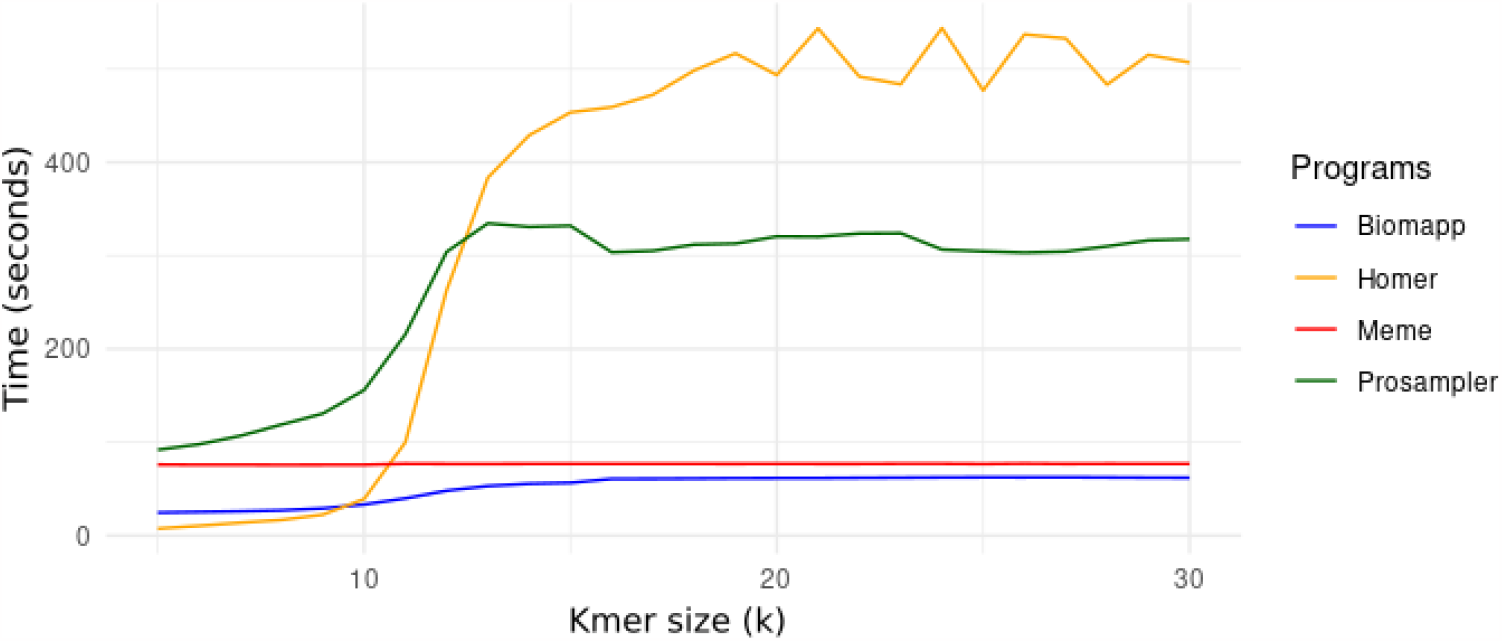
Variação do tempo de execução em relaçao a k para os algoritmos biomapp ::chip, meme, homer, e prosampler. Observa-se que o algoritmo biomapp ::chip apresenta vantagem em termos de eficiencia temporal seguido pelo algoritmo meme.

The Figure 4 complements the insights obtained in the previous figure by visualizing the time consumption of each algorithm through a bar chart. This representation makes the contrast between the time performances of the algorithms even more evident. While prosampler and homer demonstrate significantly higher time consumption, the algorithms meme and biomapp ::chip continue to exhibit better time efficiency. The main difference between the two figures lies in the type of visualization: while the previous figure used a line chart to illustrate the relationship between execution time and the size of k, Figure 4 employs a bar chart, broadening our understanding of the temporal behavior of each algorithm individually.

**Fig. 4.**
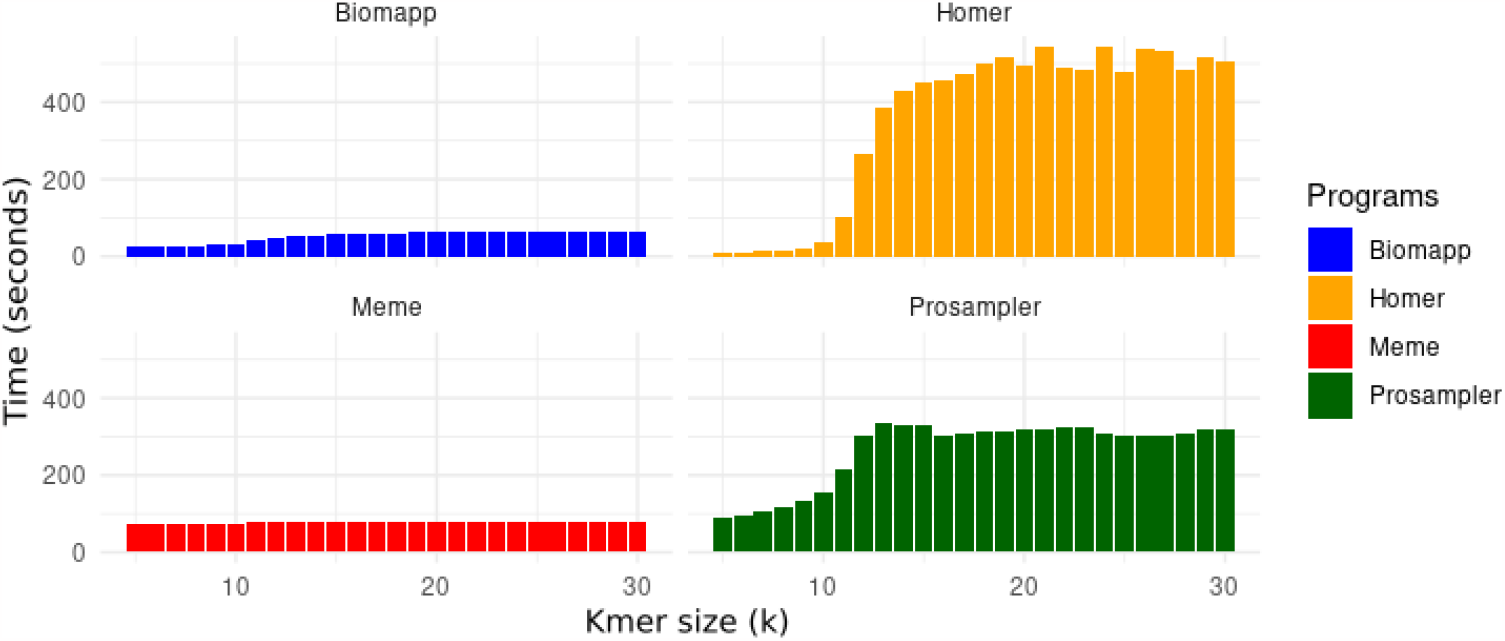
Temporal behavior as a function of the increment of k. It is noted that the execution time tends to grow with the increase of k for all the algorithms analyzed. However, such growth is more pronounced for homer and prosampler. In contrast, biomapp ::chip and meme demonstrate a more contained increase in execution time, with biomapp ::chip standing out for its better temporal efficiency.

The Figure 5 presents a line graph illustrating the RAM memory consumption for each algorithm. It is a unified graph, allowing for direct comparison between all of them. Similar to what was observed in execution time, biomapp ::chip and meme were close in memory efficiency, although meme had a slight advantage, consuming just under 500 Mbytes in the worst case. biomapp ::chip showed an average memory consumption slightly below 1000 Mbytes. In contrast, the homer and prosampler algorithms displayed much higher memory usage, reaching just over 2000 Mbytes and exceeding 4000 Mbytes in the worst scenario, respectively.

**Fig. 5.**
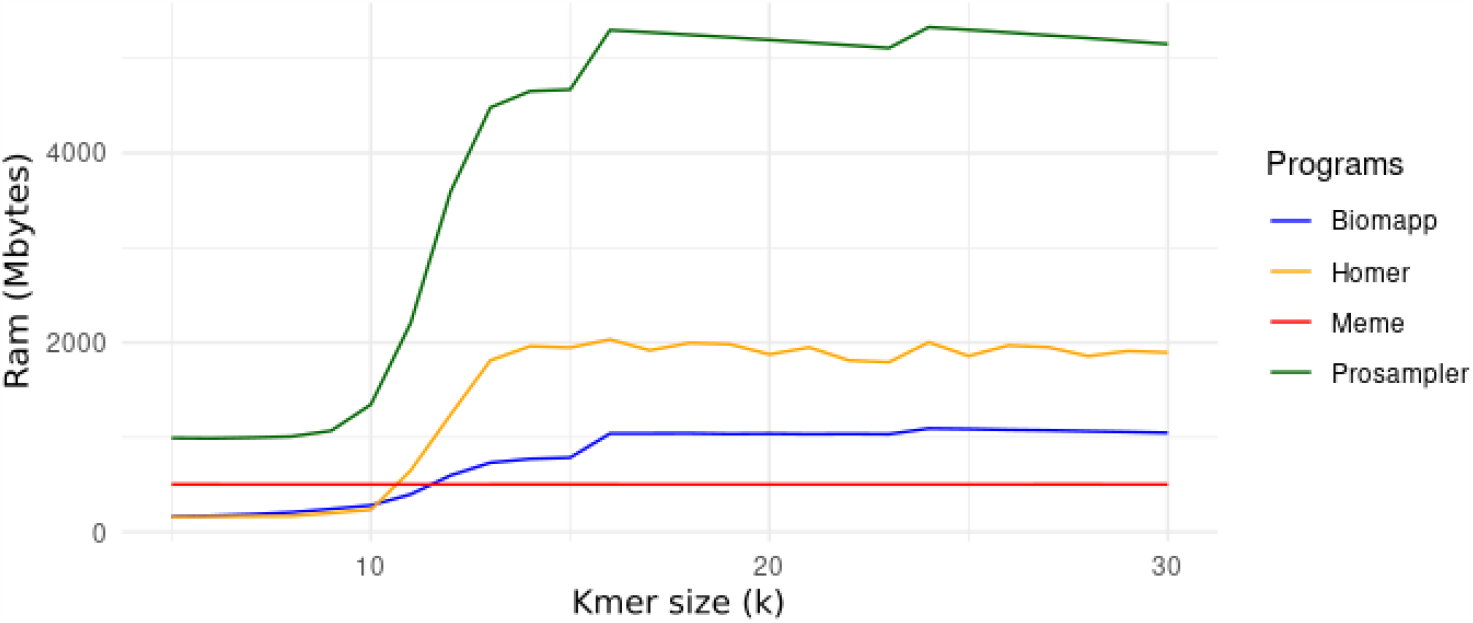
Variation of RAM memory consumption in relation to k for the algorithms biomapp ::chip, meme, homer, and prosampler. It is observed that the meme algorithm has an advantage in terms of spatial efficiency, followed by the biomapp ::chip algorithm.

The Figure 6 uses bar graphs and distributes each algorithm into a separate frame to represent RAM memory consumption. This approach facilitates the detailed visualization of the individual behavior of each algorithm. While the results remain consistent with those observed in Figure 5, the separation into distinct frames allows for more focused analysis. For example, it becomes easier to isolate cases where the memory consumption of biomapp ::chip approaches 1000 Mbytes, or when MEME keeps its consumption below 500 Mbytes. Similarly, the algorithms homer and prosampler continue to show significantly higher memory consumption, making it easier to identify the points where they exceed 2000 Mbytes and 4000 Mbytes, respectively.

**Fig. 6.**
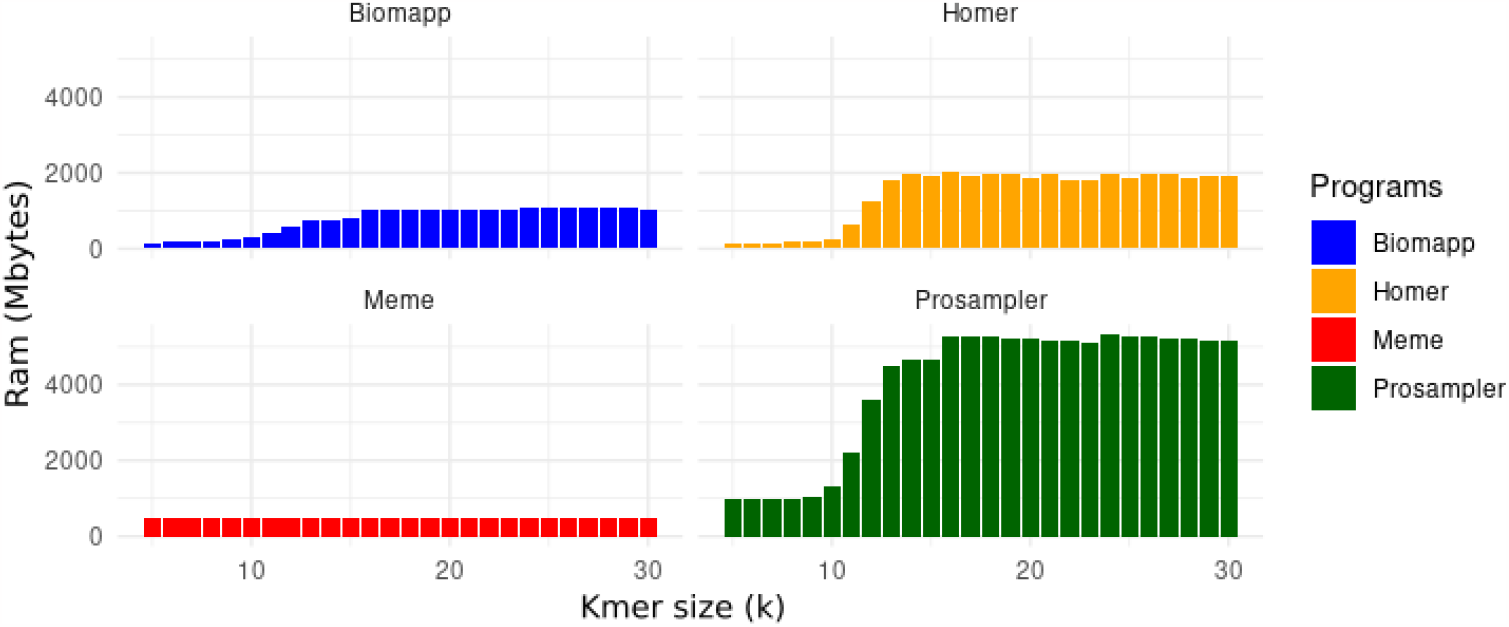
Spatial behavior as a function of the increase in k. It is noted that memory consumption tends to grow with the increase in k for all the algorithms analyzed. However, such growth is more pronounced for homer and prosampler. In contrast, biomapp ::chip and meme show a more contained increase in execution time, with meme standing out for its better spatial efficiency.

#### 3.3.2 Biomapp::chip on real data

Although experiments in simulated environments are useful for initial understanding and fine-tuning of algorithms, it is the trials with real data that provide the true validity test for any computational method in bioinformatics. They offer a much more complex and variable scenario, which more accurately plays out the conditions that algorithms will face in real-world applications. Therefore, this section is the most critical and provides more precise information about the feasibility and robustness of the evaluated algorithms.

In this test set, all algorithms were executed on a sample of 131 datasets, obtained from the jaspar repository. These data were carefully selected based on two specific criteria: i) all are from *ChIP-seq* experiments and ii) all have more than 10,000 sequences. The main focus of this test was the comparison of accuracy between the approaches. For this reason, unlike the tests performed on synthetic data, where the value of k was varied systematically, in this scenario the value of k adopted was that suggested by scientific literature. Thus, each algorithm was tested with a unique and specific value of k, according to the most recent academic guidelines.

##### Performance in relation to time and space consumption

Figure 7 reveals important nuances in the performance of the four examined algorithms in terms of execution time and ram memory consumption. biomapp:chip stands out for displaying the shortest average execution time, only 15.9 seconds, making it the most agile option. On the other hand, homer proved to be the slowest, with an average time of 624 seconds and an average ram consumption of 1436.16 MB, which could be a limitation in scenarios that demand quick responses.

**Fig. 7.**
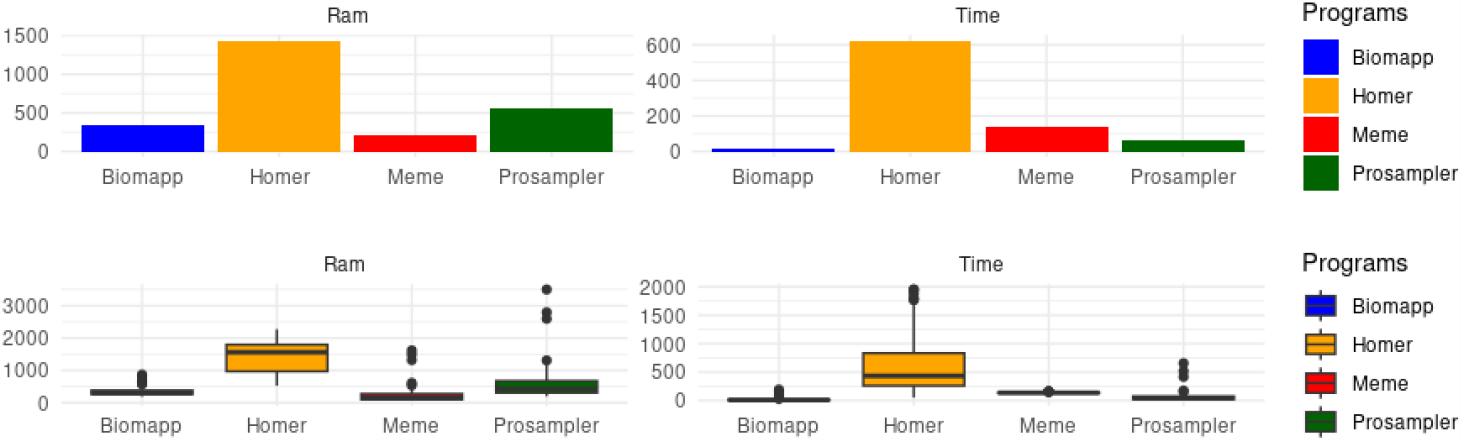
Comparison of the overall average ram memory (measured in Mbytes) and execution time consumption (measured in seconds) among different algorithms applied to real datasets. It is observed that biomapp:chip leads in memory consumption and occupies the second position in ram consumption. meme shows the lowest ram consumption, while prosampler, despite its inferior performance in synthetic tests, shows a significant improvement, coming in second place in memory consumption. The homer algorithm does not stand out in any of the criteria.

In terms of ram consumption, meme was the most efficient with an average of 217 mbytes, while HOMER exhibited the highest consumption, with 1436 mbytes. PROSAMPLER, which had moderate performance in time, registered the highest peak in ram usage, reaching 3501 mbytes. In addition, prosampler displayed the greatest variation in both time and ram consumption, suggesting that its performance may be highly sensitive to the characteristics of the analyzed dataset. Overall, the tests indicated that biomapp ::chip and meme proved to be more robust, making them more predictable in their performance, as shown in Table 3.

**Table 3.** Comparison of memory consumption (measured in Mbytes) and time (measured in seconds) between the algorithms biomapp ::chip, HOMER, meme and PROSAMPLER.

| ALGORITHM | AVG TIME | AVG RAM | MAX TIME | MAX RAM | MIN TIME | MIN RAM |
| --- | --- | --- | --- | --- | --- | --- |
| Biomapp | 15.86 | 335.57 | 192.79 | 868.92 | 1.41 | 190.08 |
| Homer | 624.45 | 1436.16 | 1952.69 | 2270.09 | 52.47 | 524.70 |
| Meme | 138.36 | 216.94 | 159.70 | 1618.04 | 118.93 | 69.32 |
| Prosampler | 61.83 | 548.17 | 655.92 | 3500.79 | 14.26 | 199.07 |

The analysis of the data presented in Figures 8 and 9 provides important insights into the performance and efficiency of the four algorithms evaluated in various scenarios, Parameterized by the value of k. First, it is possible to observe that the biomapp ::chip algorithm displays, on average, the lowest consumption of time and ram memory in almost all scenarios. Its average execution time ranges from 9.79 to 23.00 seconds, while the average ram usage lies between 251.50 and 838.70 mbytes. The algorithm is also consistent in maintaining a relatively low maximum and minimum execution time, as shown by the data in Table 4.

**Table 4.**
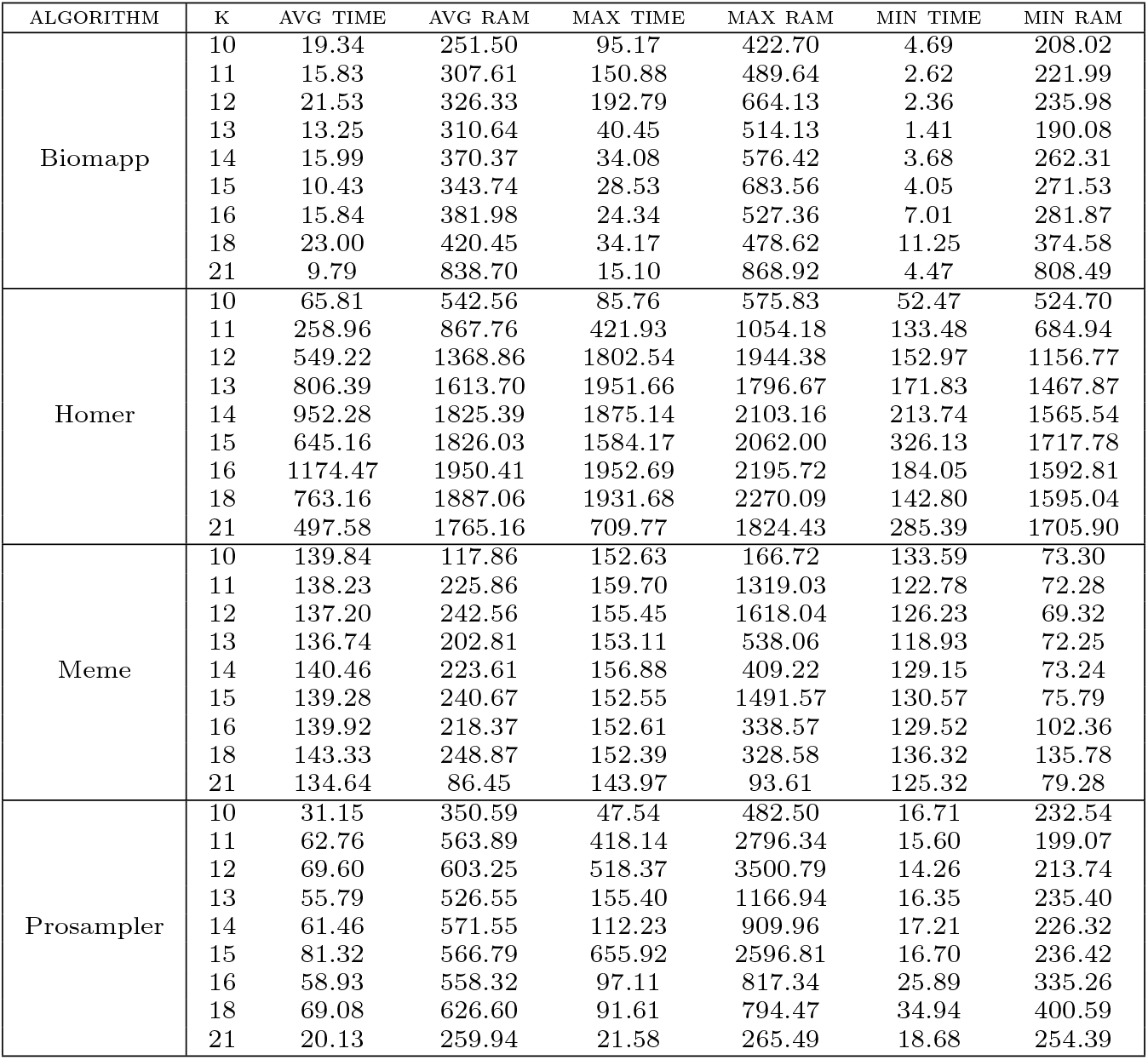
Comparison of memory consumption (measured in Mbytes) and time (measured in seconds) between the algorithms biomapp ::chip, homer, meme and prosampler grouped by k.

**Fig. 8.**
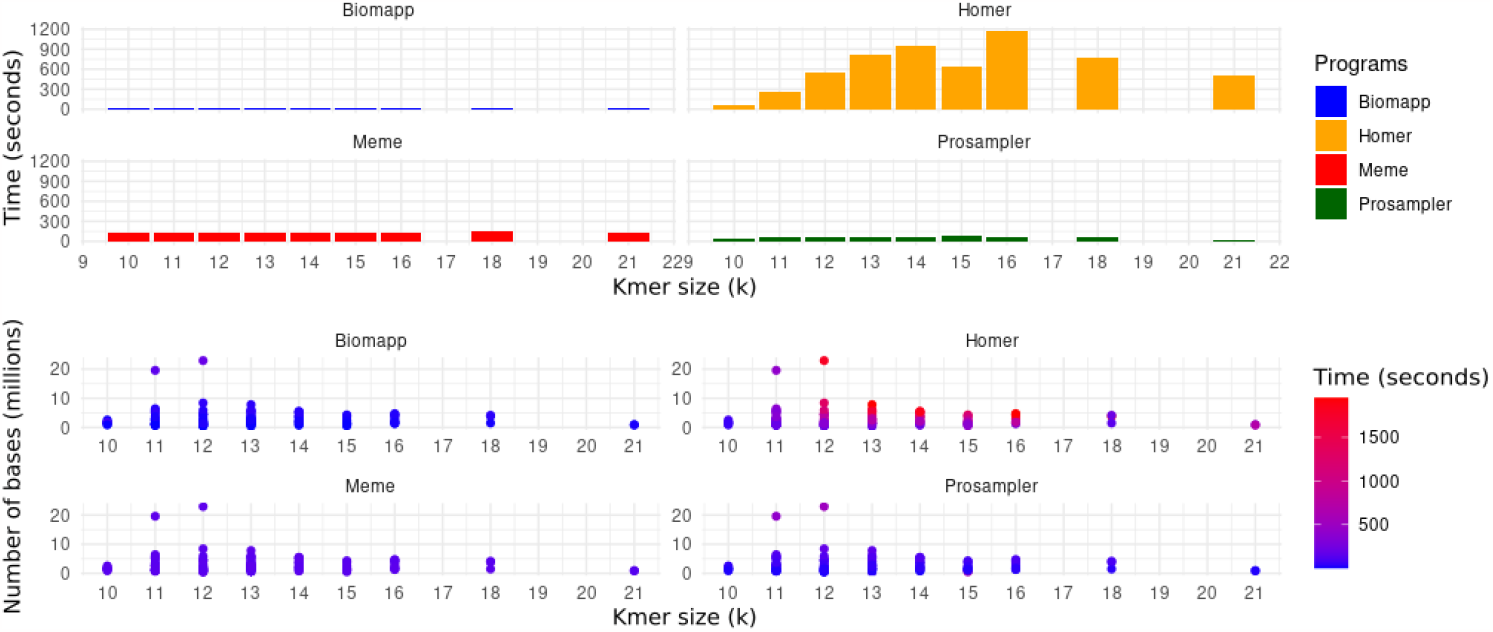
Comparison between the average performance of the algorithms in relation to time consumption grouped by value of k. It is possible to verify that biomapp ::chip, prosampler and meme exhibit the shortest times while HOMER presented significantly higher execution times.

**Fig. 9.**
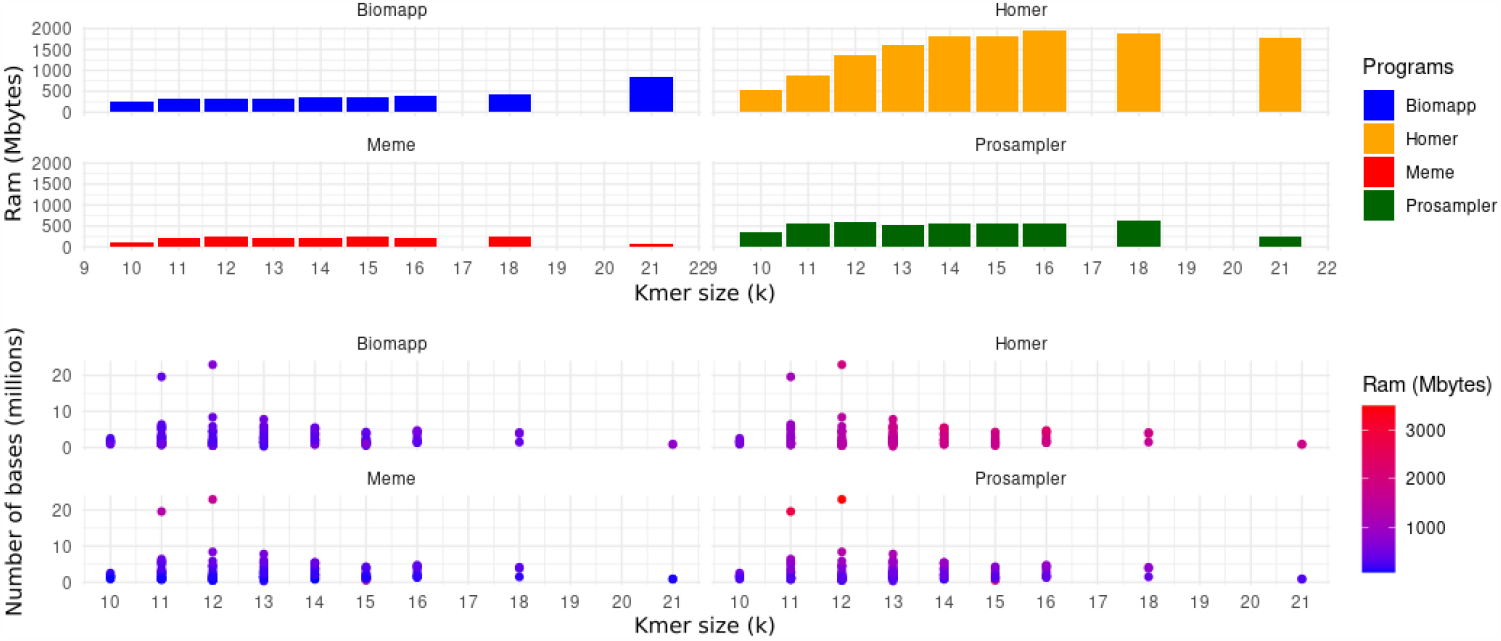
Comparison between the average performance of the algorithms in relation to memory consumption ram grouped by k value. It is possible to verify that biomapp ::chip, prosampler and meme exhibit the shortest times while HOMER presented significantly higher execution times.

On the other hand, the HOMER algorithm presents the highest consumption of both time and RAM. It is interesting to note the increase in average execution time and average ram usage as k increases, reaching a peak of 1174.47 seconds and 1950.41 mbytes for k = 16.

The meme algorithm shows relative stability in average execution time, ranging from 134.64 to 143.33 seconds, but with peaks of ram usage reaching 1618.04 mbytes for k = 12. The prosampler displays intermediate performance in terms of time efficiency and ram usage, with a large variation in maximum and minimum metrics, particularly for ram usage, which reaches up to 3500.79 mbytes for k = 12.

##### Comparison with reference models

In this subsection, we will address the evaluation of the models generated by each algorithm, contrasting them with reference models extracted from the jaspar database version 2022. The goal of this comparison is to quantify how well the algorithms were able to approximate the pwm matrices found by each approach to their respective reference matrices. For this, specific metrics were employed to measure the distance between the matrices, thus providing a comprehensive analysis of the accuracy of the methods in finding reliable and precise representations of the investigated biological motifs.

According to Figure 10 and Table 5, the biomapp algorithm showed superiority in almost all metrics analyzed. In the bha metric, the biomapp algorithm recorded an average of 0.993 and a very low standard deviation of 0.0308, signaling consistency in the results. However, it is interesting to note that prosampler had the lowest minimum in this metric (0.685), which could be a point of concern in terms of consistency.

**Table 5.**
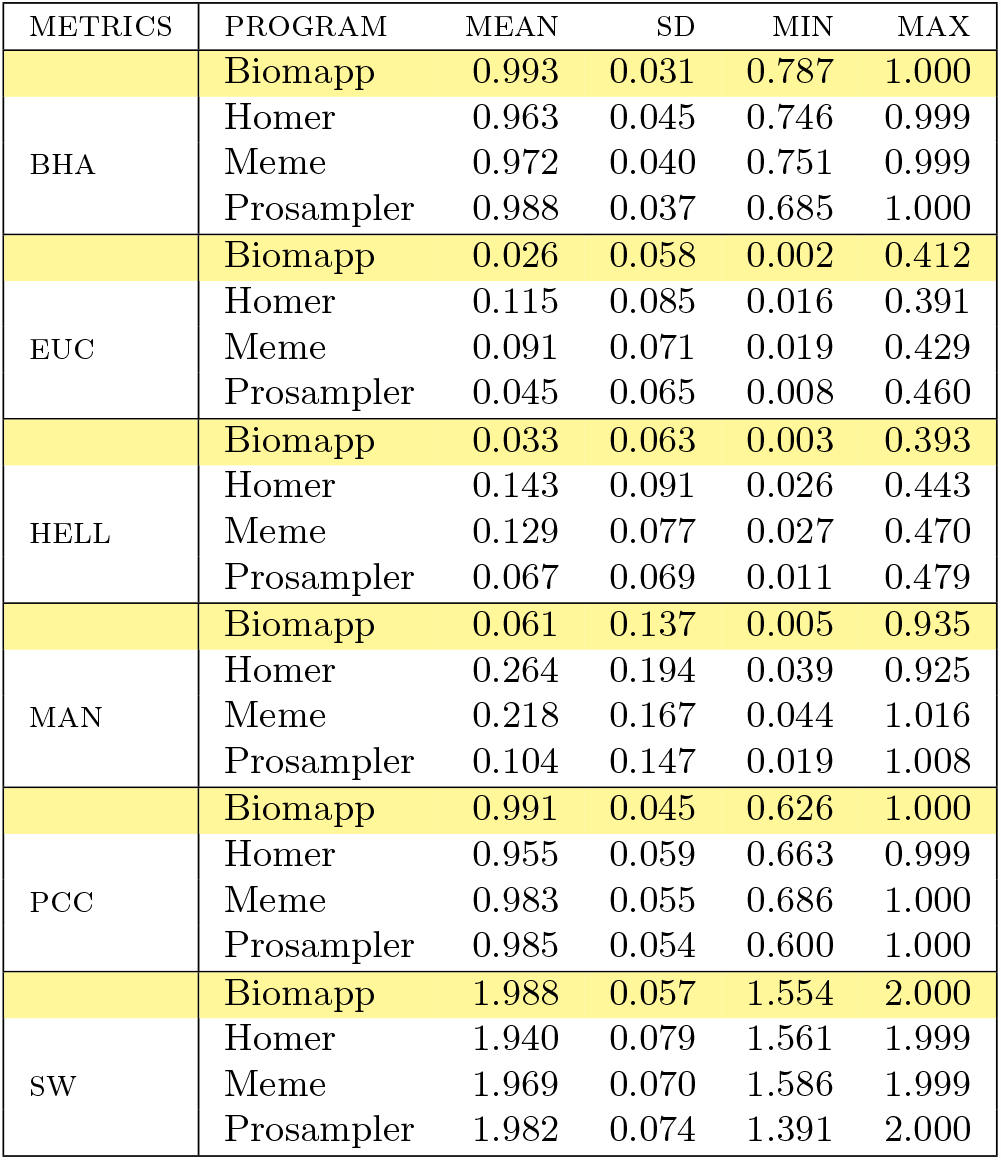
Comparative analysis between algorithms in various distance and correlation metrics. The yellow line highlights the winner of each metric in terms of average.

**Fig. 10.**
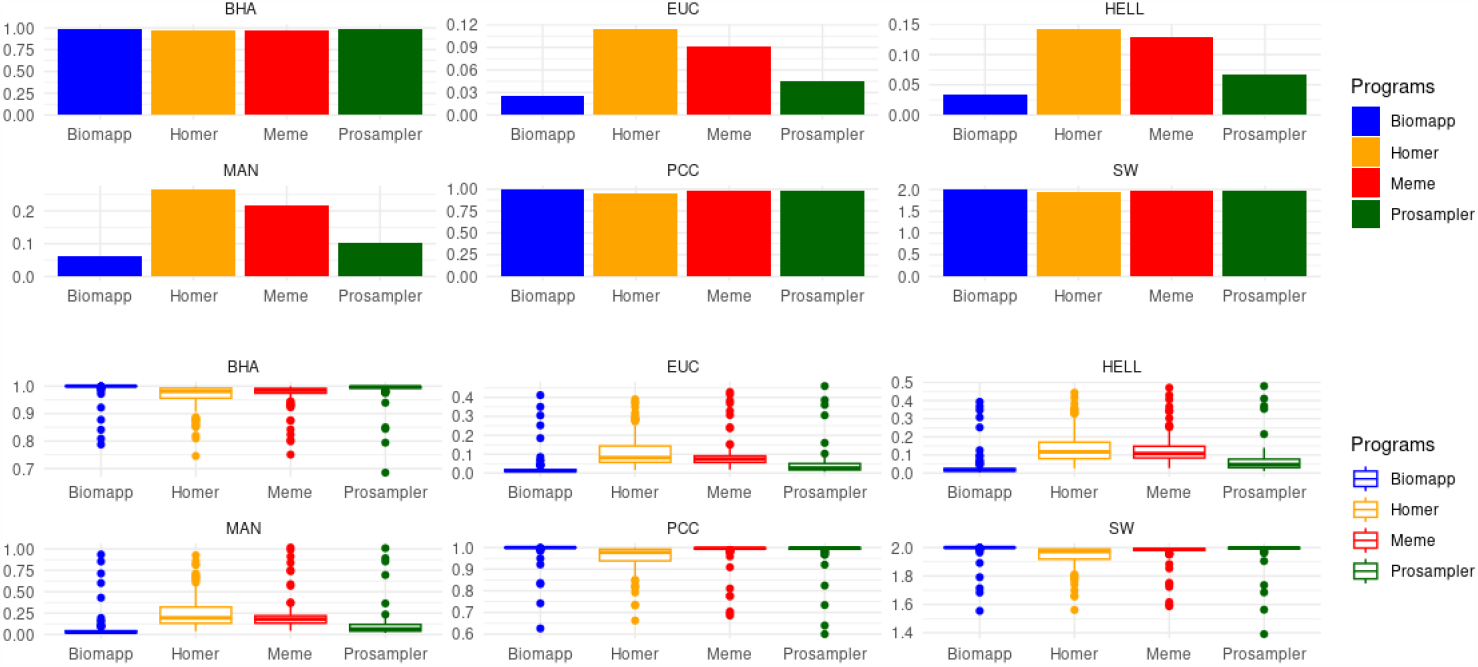
Comparative analysis of the algorithms biomapp ::chip, meme, prosampler, and homer using various distance metrics and correlation coefficients. The metrics employed for comparison include Euclidean distance, Manhattan distance, Hellinger distance, Pearson correlation coefficient, Sandelin-Wasserman coefficient, and Bhattacharyya coefficient. It is noted that, when considering the overall average, the biomapp algorithm showed superior performance compared to the other algorithms evaluated.

When we observe distance metrics such as euc, hell, and man, biomapp once again stands out for having the lowest averages. Here, homer’s performance is notably inferior; for example, in the man metric, the average is 0.264 with a standard deviation of 0.194, both values being the highest among the algorithms analyzed. This could point to a generally inferior performance of HOMER in multidimensional comparisons. Regarding the pcc metric, all algorithms showed high average values, indicating a good degree of correlation. However, it is interesting to note that prosampler had the lowest minimum value (0.600) in this metric. Such a minimum value is significantly lower than the others, which may indicate instability in specific cases. Figure 11 displays the evolution of each algorithm as a function of the variable k.

**Fig. 11.**
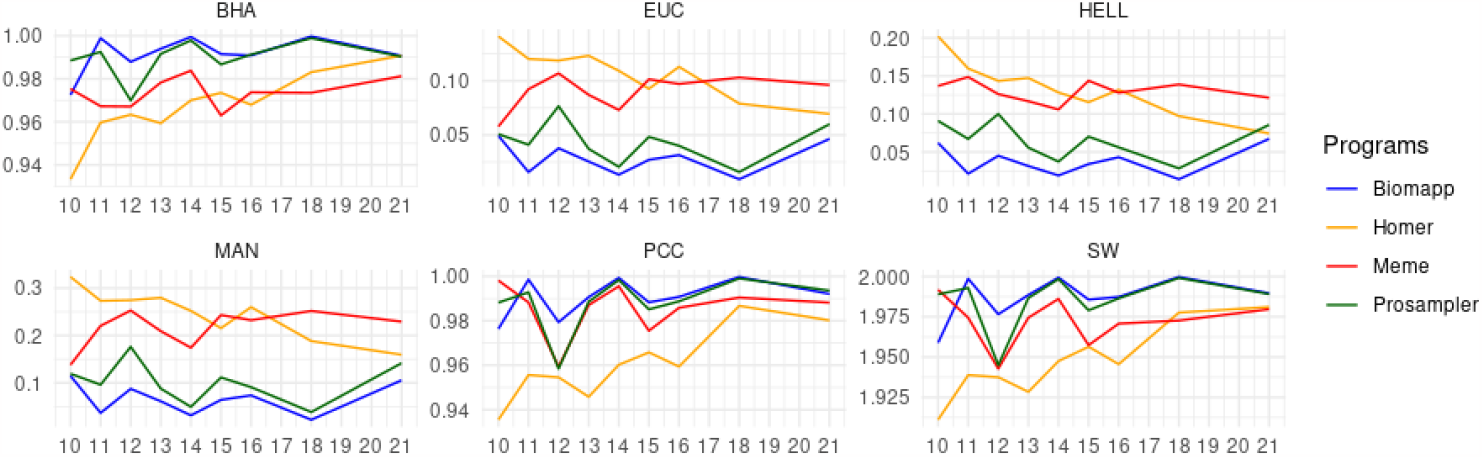
Comparative analysis of algorithms based on the evolution of metrics depending on the variable k.

It is possible to observe in this figure that BIOMAPP in the Euclidean, Hellinger, and Manhattan distances always stayed below the other algorithms, for all values of k. In the Bhattacharyya coefficient, prosampler obtained the best result (0.988) for k = 10, followed by meme (0.975) and biomapp ::chip (0.972). For other values of k, biomapp ::chip achieved better values in this metric. Something similar occurred with the Pearson correlation and Sandelin-Wasserman similarity, in which biomapp ::chip had the lowest score for k = 10.

The sw metric showed a closer competition among the algorithms, with all averages approaching 2 and relatively low standard deviations. However, prosampler here showed the lowest minimum value of 1.39, which once again raises questions about its consistency compared to the other algorithms. While biomapp exhibited a generally superior performance in all metrics evaluated, the other algorithms have weak points that deserve attention. The inferior performance of HOMER in distance metrics and the variability of prosampler in metrics like bha and pcc are points that require additional investigations.

##### Statistical analysis

In the performance analysis of algorithms, it is imperative to employ rigorous statistical techniques to determine whether the differences observed in various metrics are truly significant. The application of statistical tests provides a means to validate comparisons between different approaches or settings, ensuring that the conclusions drawn are robust and reliable.

In this subsection, use is made of the friedman Test, a non-parametric method employed to identify significant variations in medians among multiple paired groups. This statistical test is especially relevant when the conditions of normality and homoscedasticity, which are prerequisites for the application of anova, are not verified, as observed in the present study.

After the main tests, post hoc tests will be performed for multiple comparison analyses. The aim is to identify which groups are statistically different from each other. The results of the post hoc tests are essential for establishing concrete conclusions about the evaluated metrics.

The data displayed in Table 6 show the results of the friedman and NEMENYI statistical tests applied to various metrics to evaluate the performance of the motif discovery algorithms. The metrics evaluated include euc (Euclidean distance), man (Manhattan distance), pcc (Pearson correlation), hell (Hellinger distance), sw (Sandelin-Wasserman coefficient), and bha (Bhattacharyya distance).

**Table 6.** Results of friedman and nemenyi tests for the evaluation of motifs discovery algorithms. Each line represents an evaluated metric. The columns *χ*^2^ and *p−*value correspond to the results of friedman’s test. The columns relating to HOMER, meme and prosampler display the p-values obtained by the NEMENYI test, all compared to the biomapp ::chip program. The last column, called RESULT, indicates whether BIOMAPP performed better (+), lower (-) compared to the other approaches or (=) to not statistically significant.

| | $\chi^2$ | P-VALUE | HOMER | MEME | PROSAMPLER | RESULT |
| --- | --- | --- | --- | --- | --- | --- |
| EUC | 280.46 | 1.69e-60 | 0.00e+00 | 0.00e+00 | 4.01e-09 | + |
| MAN | 281.35 | 1.08e-60 | 0.00e+00 | 0.00e+00 | 8.58e-10 | + |
| PCC | 243.61 | 1.57e-52 | 0.00e+00 | 2.94e-14 | 7.42e-13 | + |
| HELL | 275.29 | 2.21e-59 | 0.00e+00 | 0.00e+00 | 1.50e-12 | + |
| SW | 261.30 | 2.35e-56 | 0.00e+00 | 0.00e+00 | 9.85e-09 | + |
| BHA | 271.19 | 1.71e-58 | 0.00e+00 | 0.00e+00 | 3.32e-11 | + |

For each metric, the friedman test produced highly significant p-values, pointing to a considerable difference between the approaches. All the p-values are very close to zero, well below the adopted significance level of *α* = 0.05, indicating that the differences are statistically significant. The winner column indicates that the BIOMAPP program outperformed all other approaches for all the tested metrics, as denoted by the symbol (+). The results of the NEMENYI test reinforce this conclusion, with p-values very close to zero. These data can be graphically visualized through Figure 12.

**Fig. 12.**
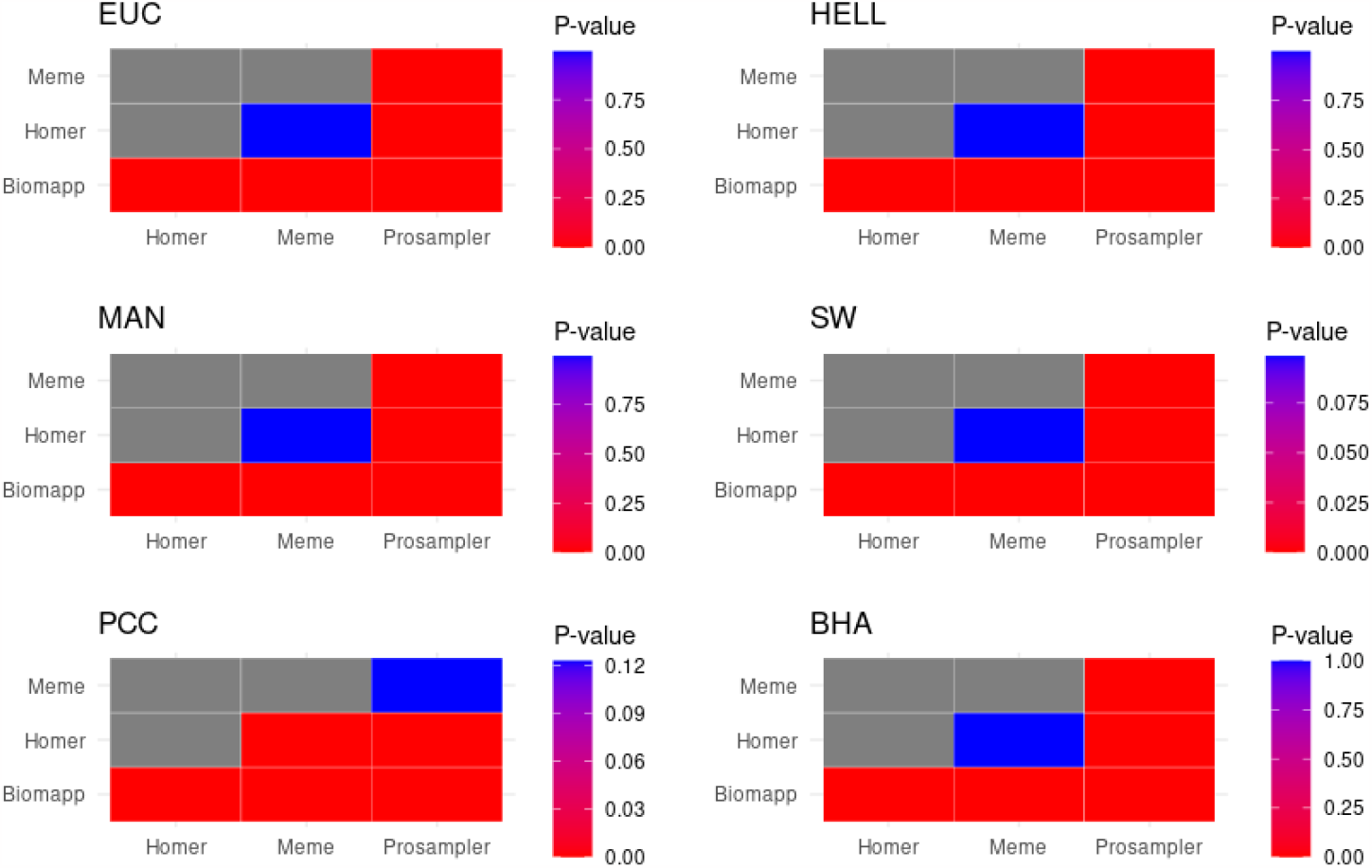
Results of p-values derived from the nemenyi test indicate the presence of significant statistical differences between the biomapp ::chip program and the other approaches tested in all metrics evaluated. Additionally, it is noted that meme does not present significant statistical differences in relation to HOMER except in the pcc metric. In the latter, both meme and prosampler were statistically similar.

Analyzing this figure, we can verify that the results obtained from the NEMENYI test point to statistically significant differences between the biomapp ::chip program and the other evaluated methodologies, covering all the metrics in question. It is relevant to highlight that meme was statistically similar to HOMER in all distance metrics. However, in the pcc metric, the meme program was more similar to PROSAMPLER, showing no statistically significant differences between them in this regard. The similarity between meme and HOMER in distance metrics and the closeness between meme and prosampler in the pcc metric can be attributed to various factors. These may include the nature of the algorithms, sensitivity to outliers, data peculiarities, or even optimization for different loss functions. Each metric may be highlighting different aspects of the relationships between the approaches, which may contribute to this behavior.

Lastly, Figure 13 presents the critical distance graphs between the different algorithms, providing an intuitive visualization of the relative positions of each method in terms of performance. The critical distance is a key value obtained from statistical tests such as the NEMENYI post-hoc test, and serves as a threshold for determining whether performance differences between multiple algorithms are statistically significant. This measure is important because it provides an objective criterion for evaluating and comparing the effectiveness of different methods. If the difference in performance between two algorithms exceeds the critical distance, we can conclude that one algorithm is significantly better than the other with a certain level of confidence.

**Fig. 13.**
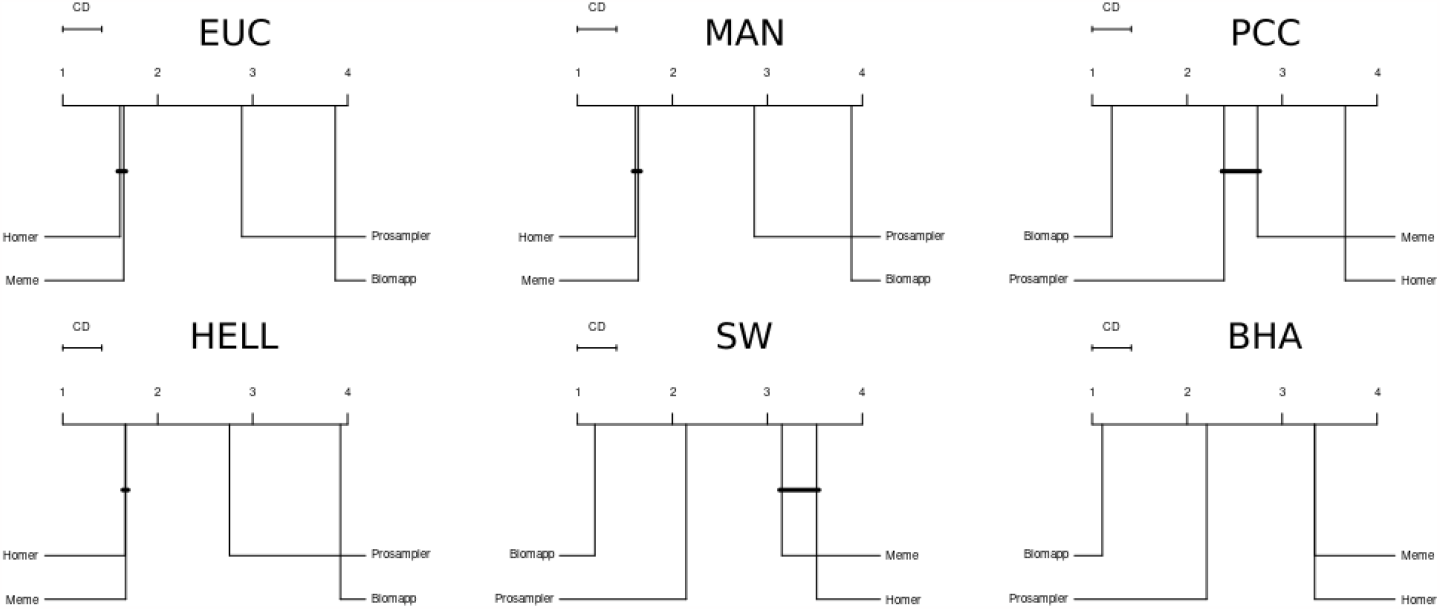
Comparison of the critical distance between different approaches. The analysis reveals that the biomapp ::chip algorithm stands out, presenting superior performance in relation to the other methodologies evaluated.

It is clearly observed that the biomapp ::chip algorithm achieved the best performance, demonstrating statistical superiority compared to all other approaches. A close relationship is also noted between the meme and homer algorithms in various metrics, especially those related to distance, corroborating the analysis performed on Figure 13. Similarly, it is also possible to see in this figure that, in the pcc metric, the meme algorithm displayed a high proximity to the prosampler algorithm, indicating that there are no statistically significant differences between them in this specific metric.

## 4 Conclusions

This study addressed the problem of biological motif discovery, a field of extreme relevance for understanding regulatory mechanisms in genomes and with implications in various areas of biology. Aiming to overcome the current limitations of available algorithms, we introduced biomapp ::chip, a tool designed to optimize both accuracy and efficiency in the analysis of large volumes of data, such as those generated by *ChIP-seq* protocols. biomapp ::chip sets itself apart with its two-step approach: the first is dedicated to efficient k-mer counting through the smt algorithm, and the second focuses on optimization via an enhanced version of the em algorithm. Our comparative analyses with state-of-the-art methods, such as meme, prosampler, and homer, demonstrated that biomapp ::chip offers a strong balance between accuracy and efficiency. The success of biomapp ::chip in the experiments suggests that the tool can be an important resource for researchers looking for reliable and effective approaches to motif discovery in large data sets. Future research may explore additional optimizations and the application of the tool in different biological scenarios.

## 5 Availability and requirements

Project name: biomapp ::chip; Project home page: https://github.com/jadermcg/BIOMAPP-CHIP; Operating system(s): Linux; Programming language: C++ and R; Other requirements: Threading Building Blocks https://www.intel.com/content/www/us/en/developer/tools/oneapi/onetbb.html; License: Apache License 2.0.

## Supplementary information

## Acknowledgments

The authors would like to thank Coordenação de Aperfeicoamento de Pessoal de Nível Superior - Brasil (CAPES) - Finance Code 001 - for the financial support given to this research.

## Declarations

### Ethics approval and consent to participate

Not applicable.

### Consent for publication

Not applicable.

## Availability of data and materials

All datasets used in the paper can be downloaded in: https://github.com/jadermcg/BIOMAPP-CHIP/tree/main/datasets.

## Competing interests

The authors declare that they have no competing interests.

## Funding

This work has been supported by Coordenação de Aperfeicoamento de Pessoal de Nível Superior - Brasil (CAPES) under Finance Code 001.

## Authors’ contributions

JMCG conceived and designed the approach. ATPR and DSS oversaw and coordinated the project. JMCG developed, implemented, realized the experiments and analyzed the results. JMCG, ATPR and DSS tested the algorithm and wrote this paper. All authors approved the final version of this manuscript.

## Footnotes

1 The resolution of the reads heavily depends on the sequencing technology employed.

## Notes

### Competing Interest Statement

The authors have declared no competing interest.

## References

[1] Altschul SF, Erickson BW (1985) Significance of nucleotide sequence alignments: a method for random sequence permutation that preserves dinucleotide and codon usage. Molecular biology and evolution 2(6):526–538

[2] Archbold J, Johnson N (1958) A construction for room’s squares and an application in experimental design. The Annals of Mathematical Statistics 29(1):219–225

[3] Bailey TL, Elkan C, et al (1994) Fitting a mixture model by expectation maximization to discover motifs in bipolymers. UCSD Technical Report CS94-351

[4] D’haeseleer P (2006) How does dna sequence motif discovery work? Nature biotechnology 24(8):959–961

[5] D’haeseleer P (2006) What are dna sequence motifs? Nature biotechnology 24(4):423–425

[6] Fitch WM (1983) Random sequences. Journal of Molecular Biology 163(2):171–176

[7] Garbelini JMC, Sanches DS, Pozo ATR (2022) Expectation maximization based algorithm applied to dna sequence motif finder. In: 2022 IEEE Congress on Evolutionary Computation (CEC), IEEE, pp 1–8

[8] Garbelini JMC, Sanches DS, Pozo ATR (2022) Towards a better understanding of heuristic approaches applied to the biological motif discovery. In: Brazilian Conference on Intelligent Systems, Springer, pp 180–194

[9] Hashim FA, Mabrouk MS, Al-Atabany W (2019) Review of different sequence motif finding algorithms. Avicenna journal of medical biotechnology 11(2):130

[10] He Y, Shen Z, Zhang Q, et al (2021) A survey on deep learning in dna/rna motif mining. Briefings in Bioinformatics 22(4):bbaa229

[11] Heinz S, Benner C, Spann N, et al (2010) Simple combinations of lineagedetermining transcription factors prime cis-regulatory elements required for macrophage and b cell identities. Molecular cell 38(4):576–589

[12] Kumar A, Hu MY, Mei Y, et al (2023) Cssq: a chip-seq signal quantifier pipeline. Frontiers in Cell and Developmental Biology 11:1167111

[13] Li Y, Ni P, Zhang S, et al (2019) Prosampler: an ultrafast and accurate motif finder in large chip-seq datasets for combinatory motif discovery. Bioinformatics 35(22):4632–4639

[14] Marçais G, Kingsford C (2011) A fast, lock-free approach for efficient parallel counting of occurrences of k-mers. Bioinformatics 27(6):764–770

[15] Norvig P, Russell S (2013) Inteligência Artificial, 3rd edn. Elsevier, USA

[16] Pevzner PA, Sze SH, et al (2000) Combinatorial approaches to finding subtle signals in dna sequences. In: ISMB, pp 269–278

[17] Sanderson C, Curtin R (2016) Armadillo: a template-based c++ library for linear algebra. Journal of Open Source Software 1(2):26

[18] Smit AF, Hubley R, Green P (1996) Repeatmasker

[19] Stormo GD (2000) Dna binding sites: representation and discovery. Bioinformatics 16(1):16–23

[20] Tatusov R, Lipman D (1996) Dust, in the ncbi. Toolkit available at http://blastwustledu/pub/dust

[21] Zang C, Schones DE, Zeng C, et al (2009) A clustering approach for identification of enriched domains from histone modification chip-seq data. Bioinformatics 25(15):1952–1958

[22] Zhang Y, Liu T, Meyer CA, et al (2008) Model-based analysis of chip-seq (macs). Genome biology 9(9):1–9

