## Supplementary file for "BIOMAPP::CHIP: Large-Scale Motif Analysis"

### Supplementary Material for BIOMAPP::CHIP: Large-Scale Motif Analysis

Jader M. Caldonazzo Garbelini

#### 1 Introduction

The purpose of this supplementary material is to provide an extended and thorough analysis focused on the comparative assessment between the SMT and JELLYFISH [1]. While the main manuscript accentuates the efficacy and efficiency of the SMT vis-à-vis other methods, with a particular emphasis on its utility in biological motif analysis, additional aspects and details warrant further investigation.

Specifically, this supplement introduces algorithms that operate over the SMT, which were not discussed in the main paper. Algorithms such as KSEARCH, IUPACSEARCH, and KHMAP will be considered herein, each with their unique peculiarities and advantages for various research scenarios in computational biology. The aim is for this material to complement and extend the findings and discussions presented in the main manuscript, thereby affording the reader a more comprehensive understanding of the versatility and applicability of the SMT in comparison to Jellyfish.

#### 2 Algorithms

In this supplementary section, we discuss algorithms that were not covered in the main article. The purpose of this additional content is to provide a comprehensive view of the methodologies we explored. Detailed pseudocode for each algorithm is also included to offer further insight into their operational mechanisms.

#### 2.1 KSearch

This algorithm's primary objective is to perform exact searches on the SMT data structure. It stands out for its efficiency, operating with a time complexity of $O(k)$, where $k$ is the size of the KMER to be found. Consequently, the algorithm traverses the fragment only once, conducting direct queries to the SMT structure to find exact matches. Its efficiency makes it particularly useful in scenarios where it is imperative to minimize search time, which is often the case in bioinformatics and computational biology applications. Algorithm 1 illustrates how this functionality was implemented.

**Algorithm 1** KSEARCH

```
k ← kmer.size()
next ← 0
for i = 1 to k do
    c ← kmer[i]
    next ← SMT(next, c)
    if next == 0 then return False
    end if
end for
return True
```

The KSEARCH algorithm is quite straightforward and starts by determining the size of the KMER, storing this information in the variable `k`. It then initializes the variable `next` with the value 0. This variable will be used to maintain the current state of the search within the SMT structure. After this, the algorithm enters the main LOOP that iterates through each character `c` of the KMER. Within this LOOP, the SMT is called with the current state `next` and the character `c` to obtain the next state. If at any point the state `next` becomes 0, the algorithm concludes that the KMER was not found and returns FALSE. The efficiency of this algorithm is highlighted by its time complexity $O(k)$, which is linear with respect to the size of the KMER. This makes KSEARCH an extremely effective tool for exact searches within the SMT.

#### 2.2 KIUPACSearch

The IUPAC-SEARCH algorithm is a specialized search function designed to identify and count patterns specified in the IUPAC format within an SMT.

The IUPAC format is a standard way of representing nucleic acid sequences, making this algorithm particularly useful for bioinformatics and the study of genetic sequences. When an IUPAC pattern is provided to the function, it traverses the SMT to identify all instances of that pattern.

The function then returns all KMERS that match the IUPAC pattern along with their counts, stored in a map where each KMER is mapped to its respective count. This provides a high-level and concise summary of all the locations in the SMT where the pattern of interest occurs, as well as the frequency with which each associated KMER is found. The IUPAC notation for nucleic acids, used for representing genomic sequences, includes letters for the four standard nucleotides and also letters to represent possible combinations thereof. Below are the definitions for IUPAC letters that represent degenerate patterns.

- R: represents a purine, i.e., an abbreviation for a position that could be either Adenine (A) or Guanine (G).
- Y: represents a pyrimidine, i.e., an abbreviation for a position that could be either Cytosine (C) or Thymine (T).
- S: STRONG or *strong interaction*, a 3-hydrogen bond. Represents a position that could be either Guanine (G) or Cytosine (C).
- W: WEAK or *weak interaction*, a 2-hydrogen bond. Represents a position that could be either Adenine (A) or Thymine (T).
- K: KETO, i.e., the nucleotides with ketone groups. Represents a position that could be either Guanine (G) or Thymine (T).
- M: AMINO, i.e., the nucleotides with amino groups. Represents a position that could be either Adenine (A) or Cytosine (C).
- B: represents a position that could be any base except Adenine (A) (i.e., could be C, G, or T).
- D: represents a position that could be any base except Cytosine (C) (i.e., could be A, G, or T).
- H: represents a position that could be any base except Guanine (G) (i.e., could be A, C, or T).

- V: represents a position that could be any base except Thymine (T) (i.e., could be A, C, or G).
- N or X: represents any base, i.e., an abbreviation for a position that could be Adenine (A), Guanine (G), Cytosine (C), or Thymine (T).

Algorithm 2 illustrates the workings of the IUPAC-SEARCH. The algorithm starts with a condition that checks if the variable $i$ is equal to $k$ in line 1. If both are equal, it indicates that the algorithm has reached the end of the IUPAC pattern it is searching for in the tree. In this case, between lines 2 to 5, the algorithm retrieves the count and the address of the current node in the tree, logs the count in the pattern map corresponding to the address, and then terminates the algorithm's execution.

If $i$ is not equal to $k$, the algorithm proceeds to get the current character from the IUPAC pattern and initializes the variables *symbol* and *next* to zero on lines 8 to 9. A new condition is checked at line 10, where if the current character is one of the nucleotide bases (A, C, G, T), the algorithm executes the instructions on lines 11 to 15. In this block, the algorithm transforms the current character into its corresponding integer value (using the *char2int* function), obtains the corresponding child node in the tree and, if such a node exists, recursively calls the IUPAC-SEARCH function to continue the search from that node.

On line 16, if the current character is $R$ (indicating purine), lines 17 to 22 are executed. Here, the algorithm checks each of the possible symbols for purine (A and G). For each symbol, it gets the corresponding child node in the tree and, if that node exists, recursively calls the IUPAC-SEARCH function.

On line 23, if the current character is $Y$ (indicating pyrimidine), the algorithm executes lines 24 to 28. This block is similar to the previous one, but it checks each of the possible symbols for pyrimidine (C and T). Finally, line 31 notes that other IUPAC letters (such as S, W, K, M, B, D, H, V, N, X) should follow a similar format to the one explained above. In the end, the algorithm stores all the KMERS and their respective counts in the *patterns* map.

The primary advantage of the IUPAC-SEARCH over the brute-force algorithm lies in its superior computational efficiency. While the brute-force method proceeds to exhaustively analyze all the KMERS in the dataset, attempting to find matches with the IUPAC pattern, the IUPAC-SEARCH, on the other hand, operates more intelligently and selectively on the SMT. This latter algorithm excels in terms of speed, as it is capable of pruning the search

**Algorithm 2** KIUPAC-SEARCH

```
if i == k then
    count ← SMT(node, 4)
    address ← SMT(node, 4 + 1)
    patterns[address] ← count
    return
end if
c ← iupac[i]
symbol ← 0
next ← 0
if c == A OR c == C OR c == G OR c == T then
    symbol ← char2int(c)
    next ← SMT(node, symbol)
    if next ≠ 0 then
        KIUPACSearch(SMT, patterns, iupac, next, i + 1, k)
    end if
else if c == R(puRine) then
    for symbol ∈ {A, G} do
        next ← SMT(node, symbol)
        if next ≠ 0 then
            KIUPACSearch(SMT, patterns, iupac, next, i + 1, k)
        end if
    end for
else if c == Y(pYrimidine) then
    for symbol ∈ {C, T} do
        next ← SMT(node, symbol)
        if next ≠ 0 then
            KIUPACSearch(SMT, patterns, iupac, next, i + 1, k)
        end if
    end for
end if
Other cases (such as S, W, K, M, B, D, H, V, N, X) follow a similar format.
```

tree. This means that nodes subordinate to patterns that do not meet the search criteria are not even considered, thus optimizing the search process.

#### 2.3 Khmap

The KHMAP algorithm employs a recursive approach to generate count hash tables from a previously built SMT. If an SMT was initially created with a `k` value equal to $\kappa$, the KHMAP is capable of extracting count tables for any value of $k$ such that $1 \leq k \leq \kappa$. The execution of this process is facilitated by Algorithms 3 and 4.

**Algorithm 3** COUNTING

```
function COUNTING(SMT, node, kmax, hmap, j)
if j == kmax then
    count = SMT(node, 4)
    hmap(kmer) += count return
end if
for i = 1 to 4 do
    next = SMT(node, i)
    if next > 0 then
        COUNTING(SMT, node, kmax, hmap, j + 1)
    end if
end for
```

The algorithm COUNT K-MERS initiates a recursive function that takes five arguments: the data structure SMT, the current node of the tree (`node`), the maximum size of the KMER (`kmax`), a hash map to store the counts (`hmap`), and a counter `j` to keep track during the recursion. The first thing the algorithm does is to check if the counter `j` is equal to `kmax`. If that is the case, it enters a conditional block to perform the counting. In this block, the algorithm retrieves the count value stored in the current node through SMT(`node`, 4). This count value is then added to the hash map `hmap` associated with the KMER under analysis. Shortly after, the function returns, ending this instance of recursion.

If `j` is not equal to `kmax`, the algorithm proceeds to a LOOP ranging from 1 to 4, which are essentially the numerical representations for the four nucleotides in a DNA sequence. For each iteration, the algorithm checks if there is a next valid node in the SMT tree. If there is, the function calls itself recursively, advancing to the next node and incrementing the counter `j`. This strategy allows the algorithm to efficiently explore the SMT tree and extract the counts of all possible KMERS of size up to `kmax`.

**Algorithm 4** HASH

```
function HASH(SMT, node, k, kmax, kmer, hmap, j)
if j == kmax then
    COUNTING(SMT, node, kmax, hmap, j) return
end if
for i = 1 to 4 do
    next = SMT(node, i)
    if next > 0 then
        HASH(SMT, next, k, kmax, kmer + char(i), hmap, j + 1)
    end if
end for
```

The algorithm HASH is a natural extension of the COUNTING algorithm and serves to create a hash table for counting *kmers*. It is also a recursive function and starts with the HASH function, which takes seven arguments: the data structure SMT, the current node in the tree (`node`), the desired KMER size (`k`), the maximum KMER size (`kmax`), the current KMER sequence, the hash map (`hmap`), and a counter `j` to control the recursion. Similar to the previous algorithm, the first step is to check if the counter `j` has reached the maximum size `k`. If true, the algorithm calls the COUNT_KMERS function to perform the counting of the KMER and store it in the hash map. At this point, the function returns, ending the current instance of recursion.

Otherwise, the algorithm enters a LOOP that iterates from 1 to 4. Each value in this iteration represents one of the four nucleotides in a DNA sequence. Within the LOOP, the algorithm checks if there is a next valid node in the SMT structure. If one exists, the HASH function calls itself recursively, advancing to the next node. In addition, it also updates the value of the KMER by appending the character corresponding to the next node and increments the counter `j`. Thus, the algorithm keeps a record of all the KMERS in a hash map, providing quick and efficient access to the data. This allows the hash table to be efficiently created, navigating through the SMT tree and collecting the necessary information.

### 3 Additional analysis: SMT vs Jellyfish

In this test set, we compare the performance of SMT with the JELLYFISH algorithm. JELLYFISH is a highly efficient K-MER counting software, commonly used in genomics and bioinformatics. This algorithm employs a HASH table to store and count K-MERS, enabling the efficient use of multiple CPU cores. With its efficient approach, JELLYFISH is capable of processing large volumes of genomic data in a considerably short amount of time when compared to other conventional methods. Moreover, the algorithm is known for its flexibility, allowing the user to adjust various parameters, such as the K-MER size and the HASH table structure. Due to these features, JELLYFISH is frequently adopted in a variety of applications, ranging from basic genomic analyses to more complex tasks like genome assembly.

In comparing SMT and JELLYFISH, we are not interested in evaluating accuracy as both are designed to provide exact counts. Therefore, accuracy is not a comparison metric between these two algorithms. Instead, we focus on efficiency in terms of runtime and RAM memory usage. To establish a rigorous comparison between SMT and JELLYFISH, experiments were conducted on synthetic and real data sets.

Execution time and memory usage were measured using the UNIX/LINUX command `/usr/bin/time -v`. This command is a reliable and widely adopted tool in the scientific community for collecting these measures, providing accurate details about resource consumption. With this set of experiments, our intention was to assess the efficiency of SMT, and through these results, contribute constructively to the field of K-MER counting algorithms and their optimizations.

For a rigorous and impartial performance evaluation between SMT and JELLYFISH, we employed a carefully planned experimentation strategy, covering a total of 200 synthetic data sets. These data sets were designed to vary in the number of sequences, scaling from 1,000 to 200,000, while maintaining a constant length of 100 bases per sequence. This experimental design allowed for the simulation of realistic usage scenarios, in addition to ensuring that the results obtained are generalizable.

It is worth noting that both JELLYFISH and SMT were run using a single THREAD throughout all the tests. Although both support *multithreading*, the decision to run them on a single THREAD was made to ensure a more transparent and fair comparison. Additionally, both were run with the command line option `-s 500M`. This setting was chosen to pre-allocate 500 megabytes of memory, a common practice to optimize performance and minimize variability in time and memory consumption results.

The algorithms were tested under these conditions, with K-MER sizes ranging between 5 and 30 bases, totaling 10,400 experiments, 5,200 for each algorithm. The range of 5 to 30 bases was chosen with the aim of providing a comprehensive view of the algorithms' behavior in different scenarios. The metrics collected during these experiments encompassed both runtime and RAM memory allocation, contributing to a more complete understanding of the computational costs associated with each method. Table 1 shows the performance of SMT and JELLYFISH in this experiment.

Analyzing the contents of Table 1, we can observe that the results show trends that are important for understanding the performance of each algorithm. Initially, in terms of execution time, it is observed that the JELLYFISH algorithm has a significant advantage over SMT for small values of `k`. However, this trend reverses at a critical point identified at `k = 15`, marked on the table by the yellow band. From this point, SMT begins to exhibit superior time performance, suggesting increasing efficiency as the complexity of the task increases, particularly for larger values of `k`. This can be better observed in Figure 1, which illustrates the behavior of the experiments in relation to time consumption, space, and base count.

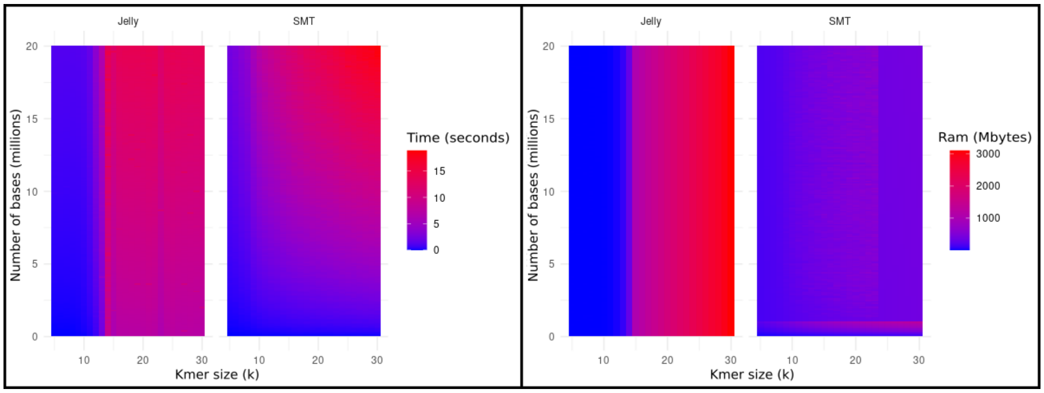


Figure 1: Heatmap illustrating the interrelationship between the parameters `k`, base count, and both execution time and RAM memory consumption, in the comparative evaluation between SMT and JELLYFISH on synthetic data.

According to these figures, for values of `k` between 5 and 15, both algorithms show highly efficient performance, represented by the blue colors on

Table 1: Comparative analysis of runtime performance and RAM memory consumption between SMT and JELLYFISH on synthetic data. The yellow band marks the transition point where SMT begins to outperform JELLYFISH.

| K | TIME JELLYFISH | TIME SMT | RAM JELLYFISH | RAM SMT |
|---|---|---|---|---|
| 5 | 0.58 | 1.18 | 4.99 | 161.27 |
| 6 | 0.58 | 1.38 | 4.99 | 171.98 |
| 7 | 0.57 | 1.69 | 4.99 | 184.79 |
| 8 | 0.58 | 2.00 | 5.06 | 203.74 |
| 9 | 0.60 | 2.91 | 5.05 | 237.04 |
| 10 | 0.70 | 3.64 | 5.76 | 276.07 |
| 11 | 1.09 | 4.14 | 9.35 | 305.74 |
| 12 | 2.37 | 4.57 | 23.31 | 332.40 |
| 13 | 4.57 | 4.90 | 79.52 | 357.26 |
| 14 | 11.35 | 5.40 | 304.30 | 377.88 |
| 15 | 9.12 | 5.77 | 1054.00 | 399.58 |
| 16 | 10.48 | 6.10 | 1203.76 | 418.64 |
| 17 | 10.60 | 6.39 | 1347.48 | 437.67 |
| 18 | 10.67 | 6.72 | 1473.34 | 456.19 |
| 19 | 10.68 | 6.95 | 1618.48 | 478.58 |
| 20 | 10.68 | 7.19 | 1752.98 | 490.08 |
| 21 | 10.89 | 7.38 | 1885.53 | 505.73 |
| 22 | 10.73 | 7.62 | 2024.81 | 520.25 |
| 23 | 9.43 | 7.85 | 2102.55 | 536.64 |
| 24 | 10.56 | 8.23 | 2293.11 | 374.75 |
| 25 | 10.75 | 8.47 | 2425.04 | 381.50 |
| 26 | 10.55 | 8.66 | 2558.47 | 387.92 |
| 27 | 10.55 | 8.86 | 2701.70 | 393.94 |
| 28 | 10.51 | 9.07 | 2801.62 | 399.64 |
| 29 | 10.49 | 9.20 | 2965.94 | 404.81 |
| 30 | 10.50 | 9.31 | 3105.55 | 409.82 |

their respective maps. However, critical differences begin to emerge as the `k` parameter increases in value. In the JELLYFISH algorithm, the `k` range from 15 to 22 reveals a transition to pinkish shades that intensify with the increasing number of bases, indicating moderate computational cost. For `k` greater than 22, the shades become even more intense, reaching nuances close to red, signaling a high computational cost.

On the other hand, the SMT algorithm reveals slightly different behavior.

For `k` greater than 12, the map remains blue up to 5 million bases but then shows a color gradient from light pink to almost red, indicating that the computational cost increases considerably for large datasets and values of `k`. This behavior was already expected for both algorithms, as the value of `k` increases, so should the computational cost related to time.

Regarding RAM memory consumption, the SMT algorithm initially requires a considerably larger volume compared to JELLYFISH. However, a phenomenon occurs again at the critical point of `k = 15`. The memory consumption by JELLYFISH increases abruptly, making SMT more efficient also in terms of memory usage.

The heat map of SMT stands out for its uniformity in the blue shade, a clear indication of more efficient computational performance in terms of RAM memory consumption. This behavior suggests a more stable and predictable resource allocation, independent of variations in the `k` parameters and the number of bases.

In contrast, the JELLYFISH algorithm exhibits more pronounced complexity. Vertical bands of colors on the map, transitioning from blue to pink and culminating in red, reveal an exponential increase in RAM memory consumption as the value of `k` grows. This characteristic implies a more sensitive and potentially limiting computational scalability, especially for large data sets and high values of `k`.

It is worth noting the scalable behavior of both algorithms. Although both SMT and JELLYFISH show an increase in time and memory metrics with the increment of `k`, SMT seems to scale in a more controlled manner. This aspect is especially observed after the transition point at `k = 15`, where SMT begins to show higher performance in both evaluated parameters.

Graphically, we can verify through Figures 2 the behavior of these algorithms in relation to time and space consumption. The SMT algorithm shows a significant performance improvement from the value `k = 15`. This improvement is particularly observable in both execution time and RAM memory usage. A careful analysis suggests that the sparse structure employed by the SMT algorithm is crucial for this performance. The use of sparse structures allows the algorithm more efficient manipulation of higher values of `k`.

In addition to the sparse structure, another factor contributing to SMT's performance is its adaptive approach to divide the dataset into batches. The batch size (`bsize`) and the number of batches (`nb`) are determined based on the relation $n \times t > 1 \times 10^6$, where `n` is the number of sequences, `t` is the

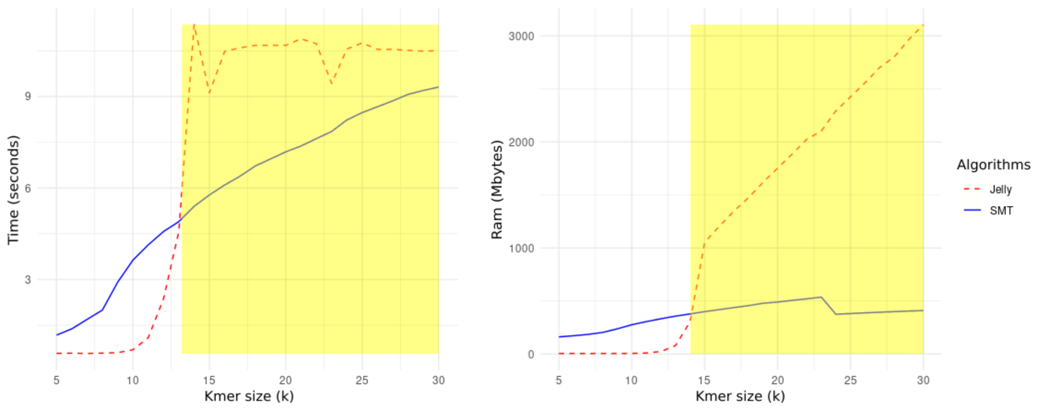


Figure 2: Comparison of time and RAM memory consumption between SMT and JELLYFISH grouped by `k` values in synthetic data. This graph provides a visual analysis of the relative performances of each algorithm for different sizes of K-MERS.

sequence width, and `k` is the size of the K-MER.

This strategy allows SMT to dynamically adjust the number and size of batches according to the specific needs of the dataset. In this way, the algorithm can optimize the use of computational resources, especially for datasets characterized by higher values of `k`, `t`, and `n`. Therefore, the transition point at `k = 15` can be justified by the interaction of these factors, highlighting the adaptability and efficiency of SMT in scenarios of high computational complexity.

To complete, we employed hypothesis tests to evaluate the performance differences between the SMT and JELLYFISH algorithms, specifically in groups segmented by different K-MER sizes. For this, we used the WILCOXON test with BONFERRONI correction. The WILCOXON test is a non-parametric test used to compare two samples and determine if their distributions are significantly different. The BONFERRONI correction, in turn, is a technique used to adjust significance levels when multiple statistical tests are performed, thereby reducing the risk of Type I errors (false positives).

The application of this correction is especially necessary in our context, as we perform multiple hypothesis tests to evaluate different K-MER sizes. The combination of these methodological approaches allowed us to perform a reliable analysis, facilitating the identification of which algorithms are more efficient in certain scenarios, thus giving greater assertiveness to our conclusions. Table 2 shows the p-values obtained with the comparison of the execution times for each algorithm.

Table 2: Statistical significance values obtained in the comparative analysis of execution time between SMT and JELLYFISH on synthetic data. The used symbology is as follows: (+) denotes superior performance of SMT, (-) points to better performance by JELLYFISH, and (=) suggests that the observed difference was not statistically significant, i.e., the null hypothesis was not rejected.

| K | JELLYFISH | SMT | W | P-VALUE | CORRECTED P-VALUE | RESULT |
|---|---|---|---|---|---|---|
| 5 | 0.57 | 1.17 | 0.00 | 3.08e-34 | 8.00e-33 | - |
| 6 | 0.59 | 1.36 | 0.00 | 1.44e-34 | 3.76e-33 | - |
| 7 | 0.57 | 1.69 | 0.00 | 1.45e-34 | 3.76e-33 | - |
| 8 | 0.58 | 2.00 | 0.00 | 2.11e-34 | 5.49e-33 | - |
| 9 | 0.59 | 2.88 | 0.00 | 1.45e-34 | 3.76e-33 | - |
| 10 | 0.69 | 3.69 | 2.00 | 1.49e-34 | 3.88e-33 | - |
| 11 | 1.13 | 4.16 | 31.50 | 2.32e-34 | 6.04e-33 | - |
| 12 | 2.49 | 4.62 | 577.50 | 6.77e-31 | 1.76e-29 | - |
| 13 | 4.62 | 4.94 | 7870.00 | 7.83e-03 | 2.04e-01 | - |
| 14 | 11.33 | 5.46 | 20100.00 | 1.45e-34 | 3.76e-33 | + |
| 15 | 9.09 | 5.89 | 20100.00 | 1.45e-34 | 3.76e-33 | + |
| 16 | 10.52 | 6.16 | 20100.00 | 1.45e-34 | 3.76e-33 | + |
| 17 | 10.61 | 6.38 | 20100.00 | 1.45e-34 | 3.76e-33 | + |
| 18 | 10.65 | 6.74 | 20100.00 | 1.45e-34 | 3.76e-33 | + |
| 19 | 10.65 | 6.97 | 20088.50 | 1.72e-34 | 4.47e-33 | + |
| 20 | 10.70 | 7.22 | 19965.50 | 1.08e-33 | 2.81e-32 | + |
| 21 | 10.89 | 7.45 | 19885.00 | 3.56e-33 | 9.26e-32 | + |
| 22 | 10.73 | 7.60 | 19327.00 | 1.06e-29 | 2.75e-28 | + |
| 23 | 9.48 | 7.82 | 15406.50 | 1.98e-11 | 5.15e-10 | + |
| 24 | 10.59 | 8.25 | 17306.00 | 8.52e-19 | 2.22e-17 | + |
| 25 | 10.78 | 8.42 | 16896.50 | 6.64e-17 | 1.73e-15 | + |
| 26 | 10.50 | 8.68 | 15716.00 | 1.36e-12 | 3.54e-11 | + |
| 27 | 10.54 | 8.80 | 15184.00 | 3.76e-10 | 9.77e-09 | + |
| 28 | 10.55 | 9.07 | 14371.00 | 1.35e-07 | 3.51e-06 | + |
| 29 | 10.44 | 9.19 | 13895.50 | 2.71e-06 | 7.05e-05 | + |
| 30 | 10.50 | 9.36 | 13600.50 | 1.48e-05 | 3.85e-04 | + |

This table presents the relevant statistical results for this investigation, including the values of W (Wilcoxon's statistic), p-values, and corrected p-values. The results indicate that for lower values of `k` (from 5 to 13), the JELLYFISH algorithm showed significantly better performance in terms of execution time. This is evidenced by the extremely low values for the p-value and corrected p-value, approaching zero. In addition, the W statistic is also very low, corroborating the superiority of JELLYFISH for these `k` values.

However, starting from `k = 14`, a reversal in the trend is observed. The SMT algorithm begins to show significantly faster execution times than JELLYFISH, as indicated by the extremely low p-values. Finally, Table 3 displays the p-values obtained through the comparison of the algorithms in relation to RAM memory consumption.

Table 3: Statistical significance values obtained in the comparative analysis of RAM memory consumption between SMT and JELLYFISH in synthetic data. The symbolism used is as follows: (+) denotes superior performance of SMT, (-) points to better performance of JELLYFISH, and (=) suggests that the observed difference was not statistically significant, i.e., the null hypothesis was not rejected.

| K | JELLYFISH | SMT | W | P-VALUE | CORRECTED P-VALUE | RESULT |
|---|---|---|---|---|---|---|
| 5 | 4.99 | 162.67 | 0.00 | 1.45e-34 | 3.76e-33 | - |
| 6 | 4.99 | 172.75 | 0.00 | 1.45e-34 | 3.76e-33 | - |
| 7 | 4.99 | 185.10 | 0.00 | 1.45e-34 | 3.76e-33 | - |
| 8 | 4.99 | 204.84 | 0.00 | 1.45e-34 | 3.76e-33 | - |
| 9 | 4.99 | 239.90 | 0.00 | 1.45e-34 | 3.76e-33 | - |
| 10 | 5.76 | 281.69 | 0.00 | 1.45e-34 | 3.76e-33 | - |
| 11 | 9.34 | 311.59 | 0.00 | 1.45e-34 | 3.76e-33 | - |
| 12 | 23.30 | 335.92 | 0.00 | 1.45e-34 | 3.76e-33 | - |
| 13 | 79.49 | 360.35 | 1.00 | 1.47e-34 | 3.82e-33 | - |
| 14 | 304.26 | 381.24 | 770.00 | 1.01e-29 | 2.64e-28 | - |
| 15 | 1053.95 | 403.02 | 20100.00 | 1.45e-34 | 3.76e-33 | + |
| 16 | 1203.71 | 420.82 | 20100.00 | 1.45e-34 | 3.76e-33 | + |
| 17 | 1347.46 | 439.79 | 20100.00 | 1.45e-34 | 3.76e-33 | + |
| 18 | 1473.28 | 460.69 | 20100.00 | 1.45e-34 | 3.76e-33 | + |
| 19 | 1618.43 | 486.16 | 20100.00 | 1.45e-34 | 3.76e-33 | + |
| 20 | 1752.96 | 492.37 | 20100.00 | 1.45e-34 | 3.76e-33 | + |
| 21 | 1885.57 | 506.75 | 20100.00 | 1.45e-34 | 3.76e-33 | + |
| 22 | 2024.83 | 516.46 | 20100.00 | 1.45e-34 | 3.76e-33 | + |
| 23 | 2102.53 | 544.90 | 20100.00 | 1.45e-34 | 3.76e-33 | + |
| 24 | 2293.12 | 363.11 | 20100.00 | 1.45e-34 | 3.76e-33 | + |
| 25 | 2425.09 | 369.34 | 20100.00 | 1.45e-34 | 3.76e-33 | + |
| 26 | 2558.46 | 375.15 | 20100.00 | 1.45e-34 | 3.76e-33 | + |
| 27 | 2701.70 | 380.63 | 20100.00 | 1.45e-34 | 3.76e-33 | + |
| 28 | 2801.66 | 385.72 | 20100.00 | 1.45e-34 | 3.76e-33 | + |
| 29 | 2965.89 | 390.26 | 20100.00 | 1.45e-34 | 3.76e-33 | + |
| 30 | 3105.54 | 394.80 | 20100.00 | 1.45e-34 | 3.76e-33 | + |

Based on Table 3, the comparative analysis of RAM memory consumption between the SMT and JELLYFISH algorithms shows an interesting dynamic across variations of the `k` parameter. For values of `k` less than 15, JELLYFISH demonstrates superior performance, consuming considerably less RAM memory. This difference is corroborated by extremely low p-values, indicating that the observations are statistically significant after correction.

However, it is essential to highlight that, starting from `k = 15`, the trend reverses. The SMT algorithm begins to exhibit much more efficient memory consumption compared to JELLYFISH. Again, the statistical significance is unequivocal, as indicated by the corrected p-values. The versatility of SMT becomes evident in this range of `k`, and this efficiency in terms of memory consumption may be important in applications requiring large-scale data analyses or in environments with limited computational resources. These findings endorse the importance of opting for the SMT algorithm in specific scenarios.

During the comparative analysis of RAM memory consumption between SMT and JELLYFISH, it was noted that many statistical tests showed identical p-values. This phenomenon can be attributed to the memory allocation mode of the two algorithms. While SMT uses an adaptive allocation strategy, varying its memory consumption according to the dataset's needs, JELLYFISH maintains a static memory allocation beyond a certain point. Specifically, for values of `k` above 14, JELLYFISH consistently allocated the same amount of memory (Table 4), regardless of the dataset size. This uniform allocation may result in less variable data distributions, contributing to the repetition of the same p-values in multiple WILCOXON tests.

One of the main advantages of SMT over other methods lies in its versatility for extracting HASH tables corresponding to KMERS of size smaller or equal to a certain value of $k$. This means that when configuring SMT with a $k = 30$ parameter, for example, there is the possibility of obtaining HASH tables for any value of $k \leq 30$. Such flexibility offers a significant advantage, allowing for a more comprehensive and adaptable analysis as needed by the research.

Table 4: Comparison table of RAM memory consumption between SMT and JELLYFISH in synthetic data.

| K | AVG JELLY | AVG SMT | MIN JELLY | MIN SMT | MAX JELLY | MAX SMT |
|---|---|---|---|---|---|---|
| 5 | 4.989 | 161.269 | 4.736 | 30.464 | 5.120 | 234.284 |
| 6 | 4.986 | 171.984 | 4.864 | 34.816 | 5.120 | 276.520 |
| 7 | 4.991 | 184.785 | 4.736 | 39.680 | 5.120 | 318.508 |
| 8 | 5.059 | 203.740 | 4.736 | 45.440 | 5.376 | 361.260 |
| 9 | 5.052 | 237.039 | 4.736 | 51.456 | 5.376 | 410.664 |
| 10 | 5.759 | 276.066 | 5.632 | 56.960 | 5.888 | 469.676 |
| 11 | 9.349 | 305.742 | 9.216 | 62.208 | 9.472 | 525.996 |
| 12 | 23.310 | 332.398 | 23.168 | 67.200 | 23.552 | 577.484 |
| 13 | 79.521 | 357.261 | 79.360 | 72.064 | 79.744 | 626.856 |
| 14 | 304.301 | 377.882 | 304.128 | 76.672 | 304.384 | 674.476 |
| 15 | 1054.001 | 399.582 | 1053.824 | 81.280 | 1054.208 | 720.680 |
| 16 | 1203.756 | 418.645 | 1203.584 | 85.632 | 1203.968 | 765.736 |
| 17 | 1347.482 | 437.674 | 1347.200 | 90.112 | 1347.584 | 809.388 |
| 18 | 1473.342 | 456.193 | 1473.152 | 94.336 | 1473.536 | 851.880 |
| 19 | 1618.484 | 478.579 | 1618.304 | 98.432 | 1618.688 | 893.096 |
| 20 | 1752.984 | 490.083 | 1752.704 | 102.400 | 1753.216 | 933.056 |
| 21 | 1885.526 | 505.729 | 1885.440 | 106.240 | 1885.696 | 971.692 |
| 22 | 2024.813 | 520.249 | 2024.576 | 109.952 | 2025.088 | 1009.196 |
| 23 | 2102.552 | 536.644 | 2102.400 | 113.536 | 2102.784 | 1045.292 |
| 24 | 2293.114 | 374.751 | 2292.736 | 116.992 | 2293.248 | 1080.364 |
| 25 | 2425.040 | 381.501 | 2424.832 | 120.320 | 2425.216 | 1114.028 |
| 26 | 2558.472 | 387.921 | 2558.208 | 123.648 | 2558.592 | 1146.540 |
| 27 | 2701.699 | 393.944 | 2701.440 | 126.720 | 2701.952 | 1177.532 |
| 28 | 2801.622 | 399.641 | 2801.408 | 129.664 | 2801.792 | 1207.596 |
| 29 | 2965.936 | 404.807 | 2965.760 | 132.480 | 2966.144 | 1236.264 |
| 30 | 3105.549 | 409.821 | 3105.280 | 135.296 | 3105.792 | 1263.788 |

### References

[1] Guillaume Marçais and Carl Kingsford. “A fast, lock-free approach for efficient parallel counting of occurrences of k-mers”. In: *Bioinformatics* 27.6 (2011), pp. 764–770.
